# A Two-Step Activation Mechanism Enables Mast Cells to Differentiate their Response between Extracellular and Invasive Enterobacterial Infection

**DOI:** 10.1101/2023.08.31.555657

**Authors:** Christopher von Beek, Anna Fahlgren, Petra Geiser, Maria Letizia Di Martino, Otto Lindahl, Grisna I. Prensa, Erika Mendez-Enriquez, Jens Eriksson, Jenny Hallgren, Maria Fällman, Gunnar Pejler, Mikael E. Sellin

**Author notes:** Correspondence: G.P.; M.E.S.

## Abstract

Mast cells (MCs) localize to mucosal tissues and contribute to innate immune defenses against infection. How MCs sense, differentiate between, and respond to bacterial pathogens remains a topic of ongoing debate. Using the prototype enteropathogen *Salmonella* Typhimurium (*S*.Tm) and other closely related enterobacteria, we here demonstrate that MCs can regulate their cytokine secretion response to distinguish between extracellular and invasive bacterial infection. Tissue-invasive *S*.Tm and MCs colocalize in the *Salmonella*-infected mouse gut. Toll-like Receptor 4 (TLR4) sensing of extracellular *S*.Tm, or pure LPS, causes a slow and modest induction of MC cytokine transcripts and proteins, including IL-6, IL-13, and TNF. By contrast, type-III-secretion-system-1 (TTSS-1)-dependent *S*.Tm invasion of both mouse and human MCs triggers rapid and potent inflammatory gene expression and >100-fold elevated cytokine secretion. The *S*.Tm TTSS-1 effectors SopB, SopE, and SopE2 here elicit a second activation signal, including Akt phosphorylation downstream of effector translocation, which combines with TLR activation to promote the full-blown MC response. Supernatants from *S*.Tm-infected MCs boost macrophage survival and maturation from bone-marrow progenitors. Taken together, this study shows that MCs can differentiate between extracellular and host-cell invasive enterobacteria via a two-step activation mechanism and tune their inflammatory output accordingly.

## Introduction

Mast cells (MCs) are innate immune cells found all over the body, but particularly enriched in barrier tissues, including the skin, the lung and intestinal mucosae. In addition to their well-known involvement in allergy, MCs take part in the host response to a wide variety of infections ^1–5^. Their strategic localization makes them frequent early responders to pathogens that disrupt epithelial linings.

In analogy with other immune cells, MCs can sense bacterial pathogen-associated molecular patterns (PAMPs) via Toll-like receptors (TLRs), e.g. TLR2 and TLR4, which detect cell wall components and lipopolysaccharide (LPS), respectively ^6–9^. Other receptors present in certain MC subsets include the peptidoglycan sensor Nod1 ^10^, and the Mas-related G-protein coupled receptor X2 (MRGPRX2) that detects bacterial quorum sensing molecules ^11^. Moreover, studies of diverse infections have demonstrated that MCs can detect assault by bacterial cytolytic toxins ^12–16^. Based on the latter studies, a common theme emerges, in which MCs respond vigorously to sublytic membrane perturbation that precedes toxin-mediated lysis ^16,17^. It is unclear whether this mode of sensing also extends to other membrane-interfering events, such as the docking of bacterial secretion machineries to the plasma membrane.

Upon IgE-crosslinking of FcεRI receptors, or in response to some infectious stimuli, MCs degranulate rapidly and release pre-stored mediators including histamine, proteases, and pro-inflammatory cytokines ^18^. Activated MCs also exhibit *de novo* biosynthesis of lipid mediators secreted within minutes, as well as cytokines and chemokines reaching detectable levels within hours ^19^. The processes of MC degranulation and cytokine/chemokine production and secretion may occur in parallel, but often appear uncoupled during infection. It remains poorly understood how MCs coordinate their different modes of bacterial sensing, and tune the nature and magnitude of their response to match the stimulus.

An additional controversy concerns the capacity MCs have to internalize bacteria. It has been proposed that bacteria, in contrast to viruses, are not internalized by MCs, and that this may explain why specifically viral infection generates a potent type I interferon response in MCs ^6^. However, other studies offer contradicting results, showing that *Staphylococcus aureus* can be internalized by murine bone-marrow-derived MCs (BMMCs), human cultured MC models and nasal polyp MCs *in vivo* ^20–23^. Further, evidence exists for some degree of MC uptake/phagocytosis also of other bacteria, e.g. non-opsonized or serum-opsonized *Escherichia coli* (*E*. *coli*) ^24,25^, *Streptococcus faecium* ^26^, and *Chlamydia trachomatis* ^27^. Thus far, no consensus has emerged on how the bacterial location affects the subsequent MC response.

Enterobacterial infections of the intestine represent one of the most prevalent classes of infectious diseases, with estimates of >600 million yearly disease cases ^28^. These infections are caused by closely related gram-negative bacteria within e.g. the *Escherichia*, *Shigella*, *Yersinia* and *Salmonella* genera. *Salmonella enterica* serovar Typhimurium (*S.*Tm) is a globally significant pathogen and a model bacterium for studies of enterobacterial pathogenesis ^29^. *S.*Tm employs flagella to swim towards the intestinal epithelium and a type-three-secretion-system (TTSS-1) to translocate effectors into targeted host cells. These effectors activate multiple Rho- and Arf-GTPases and formins, and induce actin-dependent bacterial uptake ^29,30^. By this means, *S*.Tm efficiently invades intestinal epithelial cells, but also many other non-phagocytic and phagocytic cell types ^31–35^. Both *S*.Tm and the other related enterobacteria can, however, also prevail in the extracellular environment, raising the question how our immune cells react to such disparate microbial behaviors.

Here, we have explored the MC interaction with *S*.Tm and related enterobacteria across a panel of experimental models, combining bacterial genetics with readouts for MC activation. We find that TTSS-1-proficient *S*.Tm efficiently invade MCs, whereas *S*.Tm grown under non-TTSS-1-inducing conditions, or genetically deleted for TTSS-1 components, do not. Remarkably, the MCs can tune their cytokine response to accomplish slow and low-level cytokine production when detecting extracellular enterobacteria, but swift and full-blown cytokine production in response to invasive *S*.Tm strains. This can be explained by a two-step MC activation mechanism, whereby extracellular bacteria only fuel a TLR signal, while for invasive *S*.Tm this signal combines with additional signal(s), elicited by the TTSS-1 effectors SopB/SopE/SopE2 and involving Akt pathway stimulation. This illustrates how MCs can cater their cytokine secretion response to inform their surrounding about the virulence behavior of a bacterial intruder.

## Results

### Mast cells and *Salmonella* coexist in the infected murine gut

To investigate the distribution of MCs in the bacterium-infected gut, we utilized a mouse model of *Salmonella* enterocolitis ^36^. Mice were infected per oral gavage with 3.0-7.5×10^6^ colony forming units (CFU) of *S*.Tm*^wt^*(SL1344) for 48h. Toluidine blue staining of caecal tissue sections confirmed the presence of MCs in the mucosa and submucosa of uninfected controls and *S*.Tm-infected animals (**Figure 1A-B**). When co-staining for *S*.Tm LPS and avidin (for MCs), we reproduced the MC tissue distribution in uninfected controls (**Figure 1C**), and detected MCs close to *S*.Tm-infested regions of the intestinal epithelium (**Figure 1D-E, S1A**), as well as in the deeper mucosa (**Figure 1E, S1A**), of infected animals. Total number of tissue MCs showed a modest elevation upon infection (**Figure 1F**), predominantly explained by increased mucosal MC numbers (**Figure 1G, H**). Next, we quantified transcript levels for MC-specific proteases in caecal tissue (**Figure 1I**). mRNAs encoding the mucosal MC proteases Mcpt1 and Mcpt2, as well as the connective tissue MC protease Mcpt4, were relatively highly expressed, while median transcript levels for other connective-tissue MC proteases (Mcpt5, Mcpt6, Cpa3) approached the detection limit (**Figure 1I**). When comparing control and *S*.Tm-infected sample groups, a trend towards elevated *Mcpt2* levels was noted in the latter. Hence, MCs lodge within the mucosa and submucosa of intestinal tissue prior to *S*.Tm infection, and enrich further in the mucosa by 48h post-infection (p.i.). Notably, in infected mice exhibiting significant inflammation, we also detected avidin+ staining in the epithelium-proximal lumen, which at this stage harbors high densities of *S*.Tm mixed with extruded epithelial cells and transmigrated myeloid and lymphoid cells (**Figure 1J, S1A** ^37,38^). Frequently, luminal avidin+ material did not colocalize with a distinct DAPI+ nuclear morphology. This suggests that MCs come in close contact with invasive *S*.Tm in the superficial gut mucosa (distances quantified in **Figure S1B**), and may also enter into and eventually succumb in the *S*.Tm-filled lumen.

**Figure 1.**
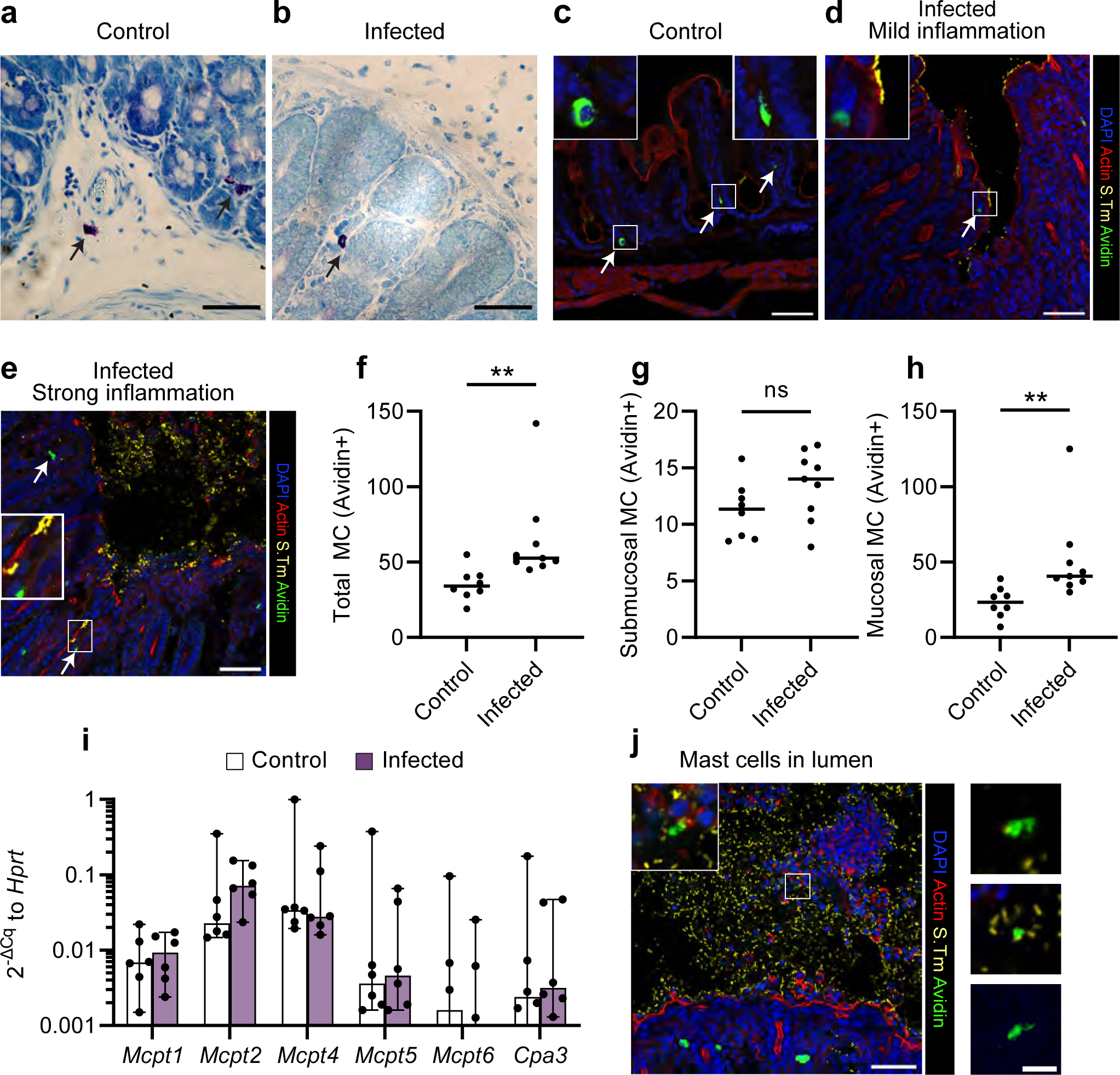
Mast cells are found in the *Salmonella*-infected caecum and come in close contact with invading bacteria. **A-B**: Toluidine blue-stained tissue sections of caecum from uninfected mice and 48h after *S.*Tm*^wt^* SL1344 infection. Arrows indicate different MC locations such as submucosa (a, bottom) and mucosa (a, top, b). Scale bars: 50µm. **C-E**: Representative IF images for caecum of uninfected mice, or infected mice in different stages of inflammation as indicated in the panel headings. Arrows indicate the position of MCs in healthy mucosa and submucosa (**C**), close to the epithelial layer (**D**) or close to invading bacteria (**E**). Scale bars: 50µm. **F-H**: Quantification of avidin+ cells per caecum section as total numbers (**F**), submucosal MCs (**G**) or mucosal MCs (**H**). Every dot indicates the mean of at least three sections for one mouse, n = 8-9. Horizontal lines display median, and significance of Mann-Whitney U test is shown. **I**: RT-qPCR analysis of total caecum tissue for mast cell protease transcripts relative to *Hprt* (2^−ΔCq^). A threshold of expression derived from Cq values < 38 was chosen. Note that some values fall under the threshold and are therefore not visible. One dot represents transcript levels in one mouse, n = 6. Bars indicate median ± 95% confidence intervals. No significance (p > 0.7) was detected for any comparison by Mann-Whitney U test. **J**: Avidin+ cells present in the lumen of infected caecum tissue sections. Representative overview image (scale bars: 50µm) and magnified images from avidin and anti-*S*.Tm-co-staining. Scale bars: 10µm.

### TTSS-1-proficient invasive *Salmonella* trigger cytokine secretion from mast cells in the absence of degranulation

To assess how MCs respond to tissue-invasive *S*.Tm, we exposed bone-marrow-derived MCs (BMMCs) to *S*.Tm*^wt^* at varying multiplicities of infection (MOI) and time frames. The *S*.Tm inoculum was cultured to promote TTSS-1 expression and invasiveness (see Methods and **Figure S2A-B**;^32,39^), similar to in the gut. BMMCs responded to the infection by secretion of IL-6 protein in a MOI- and time-dependent manner (**Figure S2C-D**). This response was most vigorous at MOI ∼25-50, and diminished again at higher MOIs (**Figure S2C**), which may be explained by dose-dependent toxicity at excessive bacterial loads. IL-6 secretion was noted from 2h p.i. and increased considerably by 3-4h p.i. (**Figure S2D**), whereas we detected significantly elevated *Il6* transcript levels already by 1h p.i. (**Figure S2E**). However, only minimal BMMC degranulation could be detected within this time frame, as assayed by β-hexosaminidase release (**Figure S2F**). We also reassessed MC degranulation using another TTSS-1-proficient *S*.Tm strain background (ATCC 14028). Again, neither *S*.Tm*^14028^ ^wt^*, nor a *S*.Tm*^14028^* ^Δ*sptP*^ mutant that lacks the SptP effector previously suggested to block MC degranulation ^40^, elicited above-background β-hexosaminidase release during the first hour (**Figure S2G**), but the calcium ionophore A23187 (positive control) did (**Figure S2F-G**). Hence, exposure to TTSS-1-primed *S*.Tm*^wt^* elicits swift *Il6* transcription and IL-6 protein secretion from BMMCs in the absence of degranulation.

Bacterial recognition by MCs has often been accredited to TLRs ^6,9^. However, work by us and others have in addition shown that sublytic levels of bacterial pore-forming toxins can elicit secretion of cytokines, including IL-6, from MCs ^14–16^. We therefore asked if MC activation by *S*.Tm could be ascribed to i) classical recognition of bacterial PAMPs, ii) membrane-perturbing effects of the TTSS-1 translocon (in analogy to pore-forming toxins), or iii) alternative mechanism(s). To address this question, we infected parallel BMMC cultures (MOI 50, 4h) with either S.Tm*^wt^*, a mutant lacking the structural TTSS-1 component InvG (S.Tm^Δ*invG*^), or one that retains the TTSS-1 apparatus and the membrane-interacting translocon, but lacks the host cell invasion effectors SipA, SopB, SopE and SopE2 (*S*.Tm^Δ4^). First, transmission electron microscopy (TEM) revealed that approximately half of all *S*.Tm*^wt^*-infected BMMCs harbored intracellular bacteria (exemplified in **Figure 2A**). The introduction of an *ssaG*-GFP reporter ^41^ (validated here against a constitutive reporter; **Figure S2B**) allowed us to assess if *S*.Tm*^wt^* and the mutant strains could invade and establish an intracellular niche within MCs. In agreement with the TEM, *S*.Tm*^wt^*efficiently invaded MCs, whereas the S.Tm^Δ*invG*^ and *S*.Tm^Δ4^ strains were non-invasive (**Figure 2B-C**). Strikingly, while *S*.Tm*^wt^* elicited prompt secretion of IL-6 (∼500pg/ml), IL-13 (∼50pg/ml), and TNF (∼50pg/ml) from the BMMCs, S.Tm^Δ*invG*^ and *S*.Tm^Δ4^ failed to do so (**Figure 2D, S2H-I**). Following a longer 24h infection, *S*.Tm^Δ*invG*^ and *S*.Tm^Δ4^ did elicit above-background levels of IL-6 secretion, but still vastly lower than the >15000pg/ml noted for *S*.Tm*^wt^* (**Figure 2E**). These findings were generalizable also to infection of BMMCs with invasive and non-invasive *S*.Tm^14028^ strains (**Figure 2F**), and to infections of murine peritoneal-cell-derived mast cells (PCMCs) (**Figure 2G**). Moreover, *S*.Tm*^wt^* induced markedly higher levels of *Il6*, *Il13*, *Tnf*, and *Nr4a3* (a MC transcription factor responsive to other stimuli; ^16,42^) transcripts than *S*.Tm^Δ*invG*^ and *S*.Tm^Δ4^, both in BMMCs (**Figure 2H**; individual data points shown in **Figure S3A-D**) and in PCMCs (**Figure 2I**, **S3G-J**). It should be noted that some other transcripts, including *Il1b* and *Nlrp3*, were upregulated by all strains (**Figure S3E-F, K-L**), suggesting the existence of several transcriptional programs that may react to different ques.

**Figure 2.**
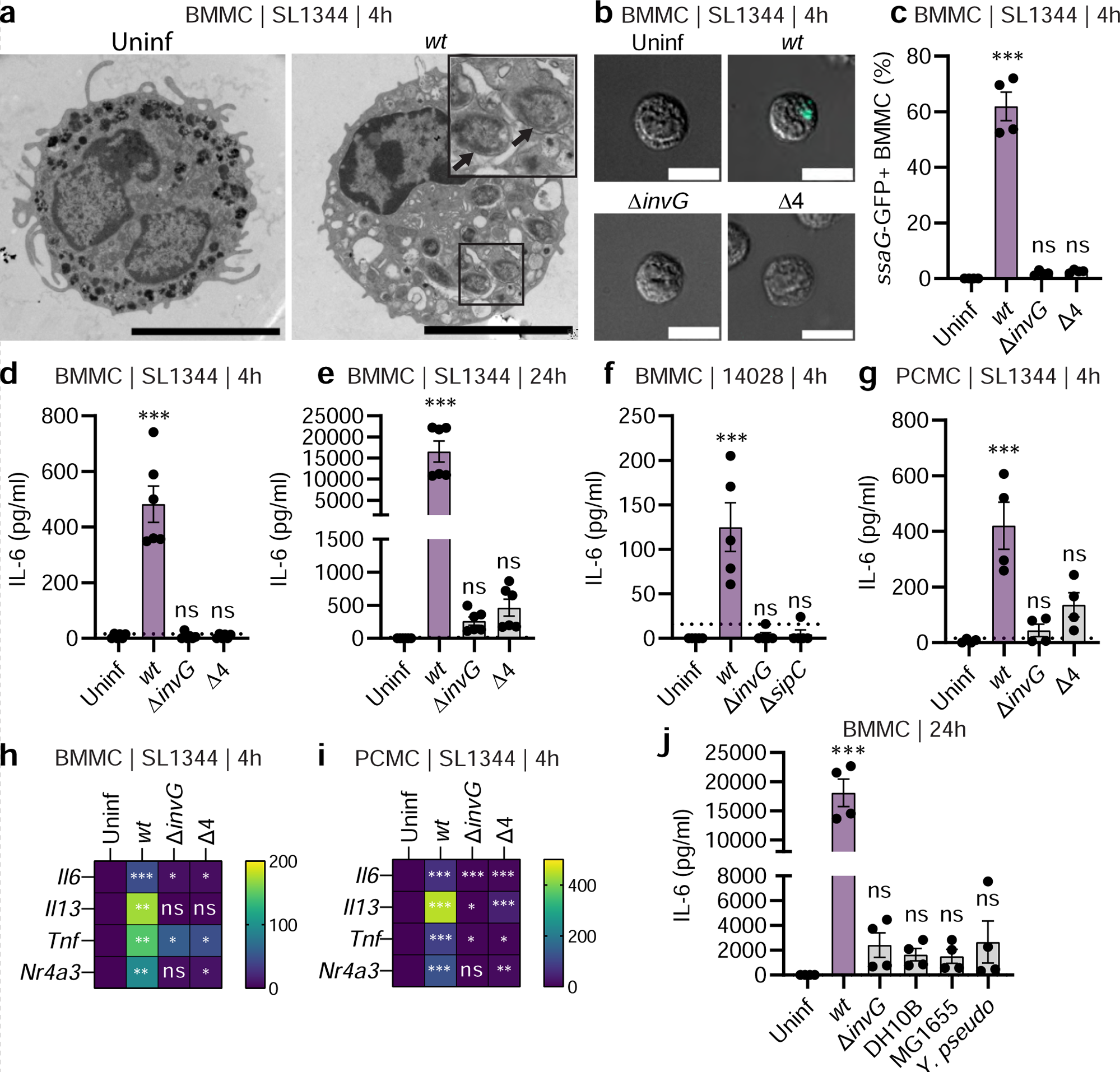
Mast cells mount a potent immunomodulatory response to *Salmonella* which is triggered by TTSS-1 effectors. **A**: Representative TEM images of BMMCs infected with *S.*Tm*^wt^* SL1344 for 4h, as well as uninfected BMMCs. Arrows indicate intracellular bacteria. Scale bar: 5µm. **B**-**C**: Representative 25 x 25µm images (**B**) and quantification by flow cytometry (**C**) of BMMCs, infected with MOI 50 of S.Tm*^wt^* SL1344 or the indicated TTSS-mutants for 4h. GFP signal and quantification show vacuolar *S.*Tm within BMMCs. **D-G** Similar conditions as above but analysis of secreted IL-6 after 4h (**D**) and 24h (**E**). **F**: Similar setup as in D, but *S.*Tm 14028 strains were used. **G:** Similar setup as in D but PCMCs were used. **H-I**: Heatmap for RT-qPCR-quantified transcript levels for *Il6*, *Il13, Tnf* and *Nr4a3* in BMMCs (**H**) and PCMCs (**I**), infected for 4h by the indicated *S*.Tm SL1344 strains. **J**: Secreted IL-6 levels from BMMCs infected with MOI 50 of S.Tm*^wt^* and *S*.Tm^Δ*invG*^ SL1344 as well as *E. coli* DH10B, *E. coli* MG1655 and *Y. pseudotuberculosis* for 24h. Every experiment was performed 2-3 times and mean ± SEM of pooled biological replicates is shown. Uninfected cells were used for statistical comparisons by ANOVA and Dunnet’s posthoc test to all other groups. For *S.*Tm ATCC 14028 infections, a Δ*malX* strain was used as *wt*.

Finally, the BMMC response to *S*.Tm was contrasted to three other non-invasive enterobacteria, namely the *E. coli* strains DH10B and MG1655, and wild-type *Yersinia pseudotuberculosis* (*Y. pseudotuberculosis^wt^*; has a TTSS apparatus, but uses it to prevent host cell uptake (^43^). Strikingly, all three *E. coli*/*Y. pseudotuberculosis* strains elicited modest levels of IL-6 secretion, and virtually undetectable IL-13 secretion, thereby phenocopying the *S*.Tm^Δ*invG*^ strain (**Figure 2J, S2J**). By sharp contrast, *S*.Tm*^wt^* again elicited a vigorous cytokine response (**Figure 2J, S2J**). We conclude that TTSS-1-proficient invasive *S*.Tm (*S*.Tm*^wt^*) trigger a potent transcriptional and cytokine secretion response in MCs, that is neither recapitulated by related non-invasive enterobacteria, nor by *S*.Tm strains genetically attenuated for invasive behavior (*S*.Tm^Δ*invG*^, *S*.Tm^Δ4^), even when these retain the membrane-interacting TTSS translocon (*S*.Tm^Δ4^, *Y. pseudotuberculosis^wt^*).

### The TTSS-1 effectors SopB, SopE, and SopE2 promote mast cell cytokine secretion upon *Salmonella* infection

Next, we surveyed the impact of individual *S*.Tm TTSS-1 effectors. Deletion of SipA (*S*.Tm^Δ*sipA*^) or SptP (*S*.Tm^Δ*sptP*^) marginally decreased, respectively marginally increased (non-significant), the levels of *S*.Tm-induced IL-6 secretion from BMMCs by 4h p.i. (**Figure 3A**). By contrast, simultaneous removal of SopB, SopE, and SopE2 (*S*.Tm^Δ*sopBEE*2^) phenocopied the *S*.Tm^Δ4^ and *S*.Tm^Δ*invG*^ strains (i.e. minimal IL-6 secretion at 4h p.i.) (**Figure 3A**). Analysis of *Il6* and *Tnf* transcripts further corroborated these findings (**Figure 3B, S4A-B**). To assess how the effectors influenced BMMC invasion, we again exploited *ssaG*-GFP reporter strains (**Figure 3C**). Flow cytometry showed that *S*.Tm*^wt^* and *S*.Tm^Δ*sipA*^ invaded BMMCs to an equal degree, while *S*.Tm^Δ*sopBEE*2^ failed to establish intracellularly, analogous to *S*.Tm^Δ*invG*^ and *S*.Tm^Δ4^ (**Figure 3C**; MOI-response curves in **Figure S4C**). Moreover, *S*.Tm^Δ*sptP*^ was hyperinvasive (**Figure 3C, S4C**). This agrees with earlier findings that this effector can counteract SopE/E2 during host cell invasion ^44^. Hence, the ability of *S*.Tm strains to trigger a cytokine response in murine BMMCs correlate with their respective ability to invade these cells.

**Figure 3.**
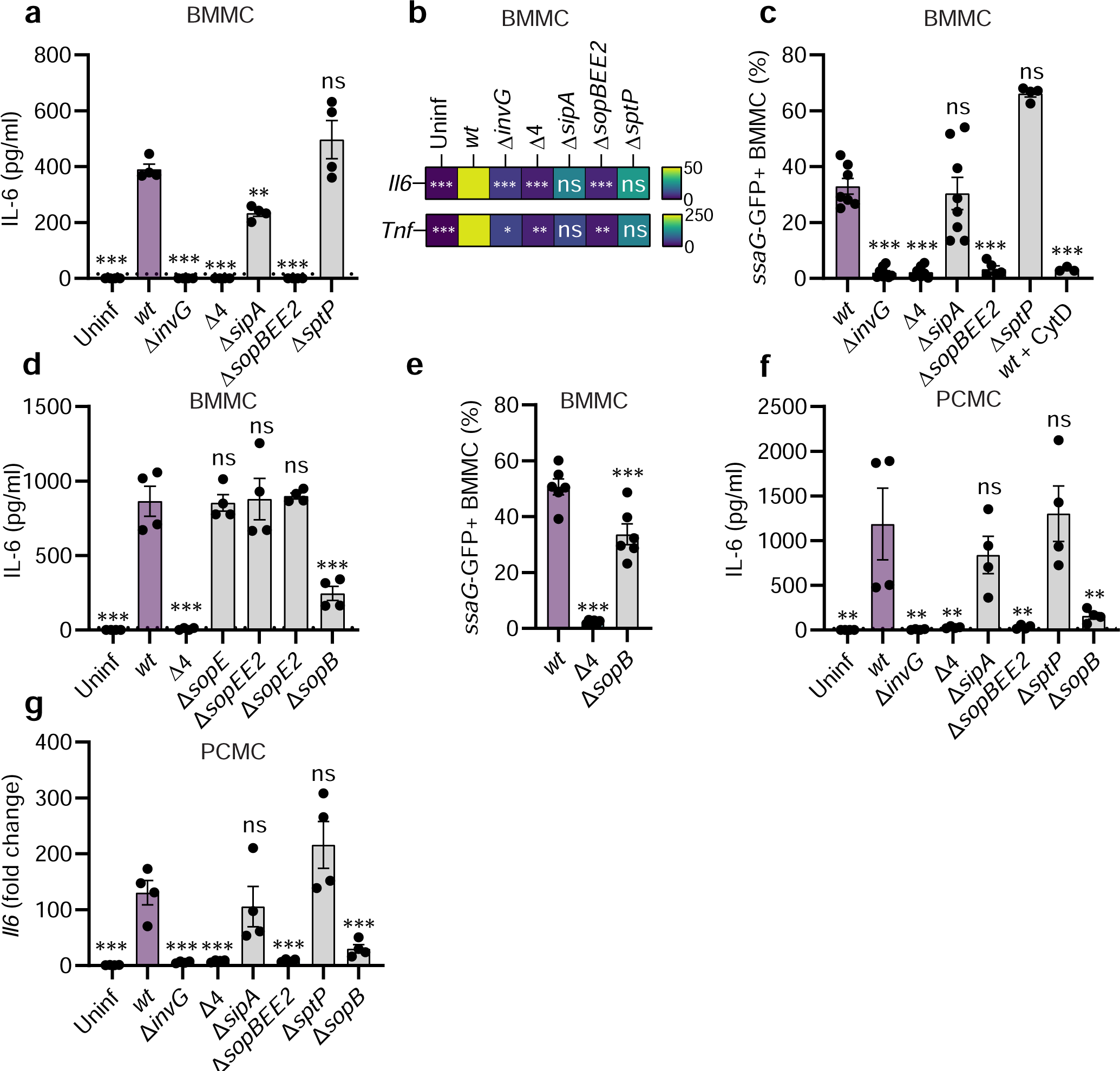
The TTSS-1 effectors SopB, SopE, and SopE2 induce mast cells cytokine expression and secretion upon *Salmonella* infection. **A-G**: BMMCs (**A**-**E**) or PCMCs (**F-G**), infected with MOI 50 of *S.*Tm*^wt^* SL1344 or the indicated TTSS-mutants of for 4h. **A, D, F** show IL-6 secretion, **B** shows a heatmap for RT-qPCR-quantified transcript levels of *Il6* and *Tnf* in BMMCs, **G** shows *Il6* transcript levels in PCMCs. **C** and **E** show percentage of BMMCs harboring vacuolar (*ssaG*-GFP+) *S.*Tm. Every experiment was performed 2-4 times and mean ± SEM of pooled biological replicates is shown. *S*.Tm*^wt^*-infected cells were used for statistical comparisons by ANOVA and Dunnet’s posthoc test to all other groups.

We next asked whether these observations generalize to human MCs, therefore turning to the untransformed MC line LUVA ^45^. LUVA cells did not secrete IL-6 (<16pg/ml), but rather appreciable amounts of TNF (∼700pg/ml) upon *S*.Tm*^wt^* infection (**Figure S4D**). Most importantly, both this cytokine response, and the capacity of *S*.Tm to invade LUVA cells, was dramatically attenuated by SopB/E/E2 deletion (**Figure S4D-E**), in full agreement with the data above.

As the *S*.Tm^Δ*sopBEE*2^ strain failed to promote invasion and cytokine secretion in murine and human MCs, the individual roles of SopB, SopE and SopE2 were explored further. Deletion of only SopE or SopE2 in isolation, or SopE/E2 in combination, had no effect on *S*.Tm’s capacity to elicit IL-6 secretion from BMMCs (**Figure 3D**). However, single deletion of SopB resulted in a drop in IL-6 secretion (**Figure 3D**). The number of BMMCs harboring intracellular (*ssaG*-GFP+) *S*.Tm was also modestly (∼2-fold) reduced by SopB deletion (**Figure 3E**; MOI-response curves in **Figure S4F**). This hints towards a role for SopB in stimulating MC cytokine secretion, and a less prominent role in driving MC invasion. Also in the PCMCs, the *S*.Tm^Δ*sopB*^ strain elicited lower IL-6 protein secretion and *Il6* transcript levels at 4hp.i. (**Figure 3F-G**). It should, however, be noted that in BMMC cultures from another wild-type mouse strain (C57BL/6J; Jackson), *S*.Tm^Δ*sopBEE*2^ again triggered negligible IL-6 secretion, while we could not substantiate a non-redundant role for SopB (**Figure S4G**). Taken together, these data demonstrate that the TTSS-1 effectors SopB/E/E2, working in partial redundancy with each other, promote a swift and full-blown MC cytokine secretion response to *S*.Tm infection.

### *Salmonella*-invaded mast cells comprise the source of cytokine production

Based on results above, it appeared likely that *S*.Tm-invaded MCs were specifically responsible for cytokine production. However, cross-talk between invaded host cells and non-infected bystanders, which respond by cytokine secretion, has been described in other contexts ^21,46,47^, and could not be ruled out. To distinguish between these possibilities, we turned to GFP-based flow cytometry sorting of BMMCs, following infection with *S*.Tm*^wt^*/*ssaG*-GFP (**Figure 4A**). The *ssaG*-GFP+ BMMC fraction expressed elevated *Il6* transcript levels, similar to the total unsorted population, and significantly higher than the *ssaG*-GFP-BMMC fraction (**Figure 4A**). The difference between *ssaG*-GFP+ and *ssaG*-GFP-fractions was, however, relatively modest (**Figure 4A**). This might be explained by either that 1) bystander BMMCs also express appreciable levels of *Il6*, or that 2) the *ssaG*-GFP-fraction contains some invaded BMMCs where the *S*.Tm had not turned on the reporter. Staining infected BMMCs for *Salmonella* LPS revealed option 2 to be true (**Figure 4B, S5A-D**; further supported by comparisons in **Figure S2B**).

**Figure 4.**
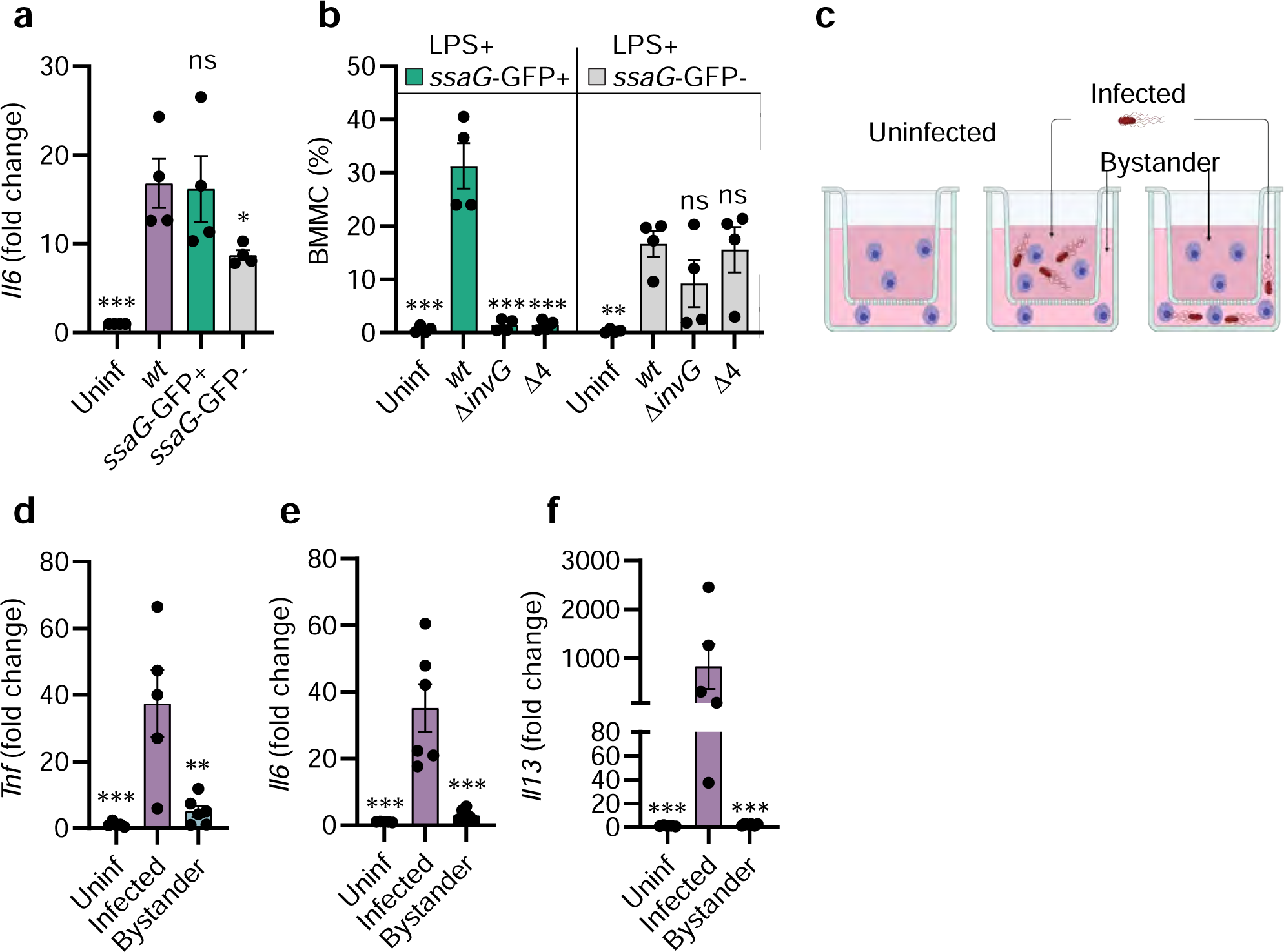
*Salmonella*-invaded mast cells are the main source of cytokines. **A:** BMMCs were infected with *S.*Tm*^wt^* SL1344 carrying the p*ssaG*-GFP reporter for 4h and sorted to enrich the *ssaG*-GFP-population. *Il6* transcript levels for both fractions as well as uninfected and unsorted *S.*Tm-infected BMMCs are shown. **B:** BMMCs were infected with *S.*Tm*^wt^* SL1344 for 4h and stained for *Salmonella* LPS. Relative population sizes of MC positive for LPS and/or vacuolar (*ssaG*-GFP+) *S.*Tm. **C**: Experimental setup for analysis of the source of soluble factors secreted by MCs **D-F**. MCs were infected with *S.*Tm*^wt^* in one compartment while separated from MCs in the other compartment not coming in direct contact with the bacteria. RT-qPCR-quantified transcript levels for *Tnf*, *Il6* and *Il13*, respectively, in the aforementioned transwell compartments. Experiments were performed 2-3 times and mean ± SEM of pooled biological replicates is shown. For A, C-F; *S*.Tm*^wt^*-infected cells were used for statistical comparison by ANOVA and Dunnet’s posthoc test to all other groups. For B, two-way ANOVA with Dunnet’s posthoc test was used to compare *S*.Tm*^wt^*-infected cells within each subpopulation to all other groups. Figure 4C was created with BioRender.com.

To separate invaded and bystander BMMCs by more definite means, we conducted infections in transwell plates with filters of 0.4 µm pore size. BMMCs were added to both top and bottom well compartments, but *S*.Tm*^wt^*was only inoculated in one of the compartments (either top or bottom; **Figure 4C**). Strikingly, BMMCs in the *S*.Tm-inoculated compartment responded with vigorous production of *Il6*, *Il13* and *Tnf* transcripts (**Figure 4D-F**). In the non-infected compartment, BMMCs can still be exposed to diffusible bacterial PAMPs, secreted BMMC factors, or DAMPs released from dying MCs, but this only led to marginally elevated cytokine transcription (**Figure 4D-F**). Transcript levels were in fact ∼7.5-fold (*Tnf*), ∼12-fold (*Il6*), to even ∼400-fold (*Il13*) higher in the *S*.Tm-infected than in the bystander compartment (**Figure 4D-F**). Hence, bacterium-invaded MCs are the main cytokine responders upon infection with TTSS-1-proficient invasive *S*.Tm.

### Combined TLR4 and SopBEE2 signals fuel cytokine secretion from *Salmonella*-infected mast cells

*S*.Tm express TLR ligands, most notably the TLR4 agonist LPS and the TLR5 agonist flagellin, which could contribute to MC activation. Measurable levels of TLR4 were detected on BMMCs (**Figure S6A**). Accordingly, BMMCs responded with IL-6 secretion upon stimulation with pure *E. coli* LPS, but not flagellin (**Figure S6B**). The LPS response was blocked by preincubating with the TLR4 inhibitor TAK242 (**Figure S6B**). Notably, the levels of IL-6 secretion elicited by LPS alone was still >25-fold lower than for a *S*.Tm*^wt^* infection (compare **Figure S6B** and **Figure 3A**). As shown in **Figure S6C**, TAK242 preincubation also attenuated the IL-6 response of BMMCs to *S*.Tm*^wt^* infection by ∼40%, without significantly affecting the number of BMMC-associated S.Tm (**Figure S6D**). Hence, TLR4 activation contributes to, but is on its own insufficient to account for the total cytokine secretion response of *S*.Tm-invaded MCs. This makes sense considering that both invasive and non-invasive S.Tm strains carry LPS, while only the former elicit full-blown MC cytokine secretion (**Figure 2-3**).

To test the relationship between *S*.Tm invasion, the TTSS-1 effectors SopB, SopE, and SopE2, and MC cytokine secretion, we next analyzed the effect of blocking bacterial internalization. Cytochalasin D (Cyto D) treatment prior to infection decreased the number of *S*.Tm*^wt^* invasion events (i.e. % *ssaG*-GFP+ BMMCs) in a dose-dependent manner (**Figure 5A**). Still, these BMMCs secreted IL-6 to a similar extent as in the absence of Cyto D pretreatment (**Figure 5B**). This suggests that while TTSS-1 and SopB/E/E2 fuel full-blown cytokine secretion (**Figure 2-3**), internalization of *S*.Tm into the MCs is not strictly required.

**Figure 5.**
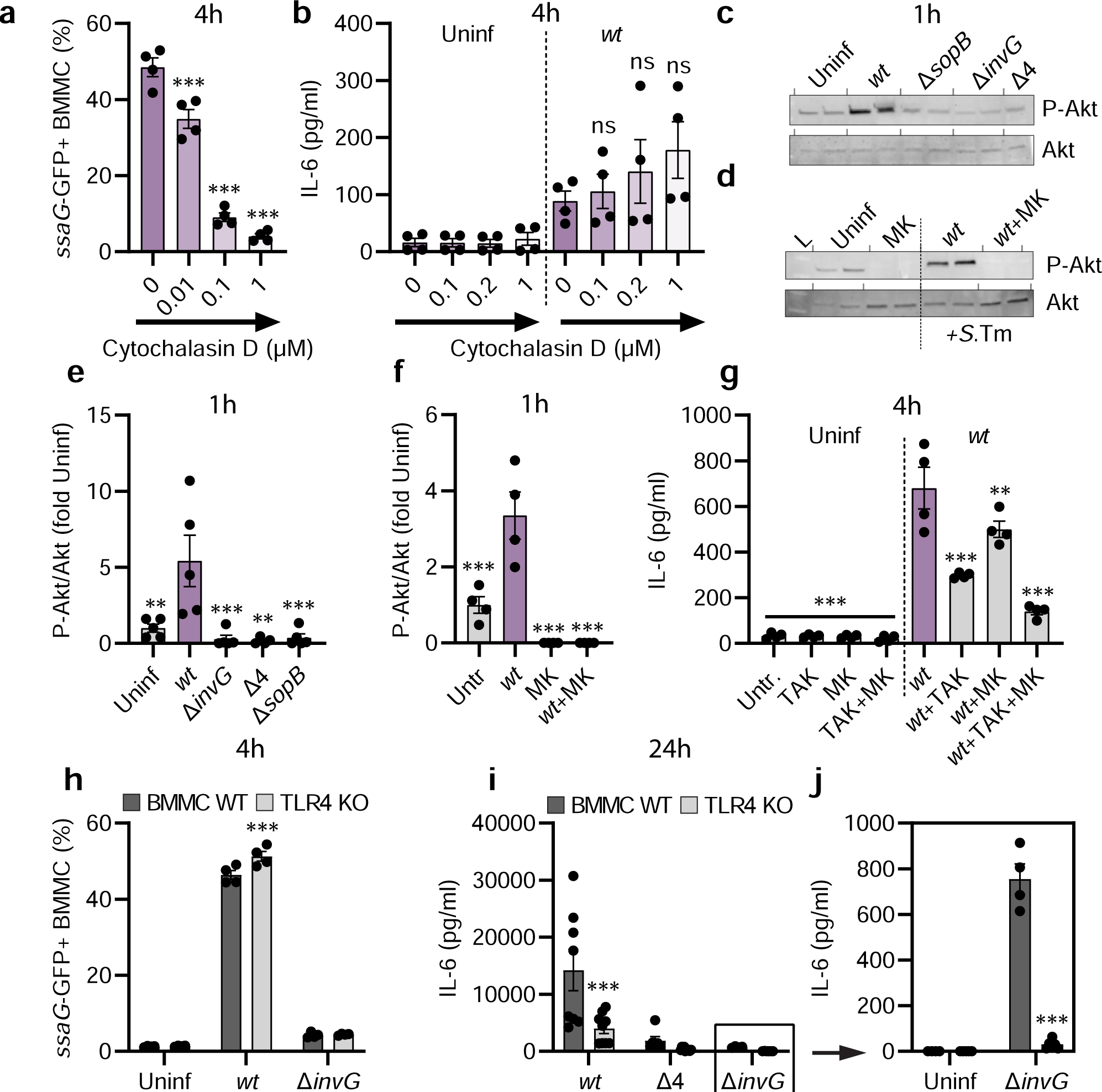
TLR4 and Sop effectors drive cytokine secretion from *Salmonella*-invaded mast cells. **A:** BMMCs were pre-treated for 1h with the Cyto D concentrations indicated, and infected with MOI 50 of *S.*Tm*^wt^* SL1344 carrying the p*ssaG*-GFP reporter. After 4h, BMMCs harboring vacuolar *S.*Tm were quantified. **B:** Similar setup as in A, but IL-6 secretion was measured. **C:** BMMCs were left uninfected or infected with MOI 50 of *S.*Tm*^wt^*SL1344 or the indicated TTSS-mutants. After 1h, cells were harvested and analyzed by immunoblot for P-Akt and Akt. **D**: BMMCs were pre-treated with MK-2206 for 1h, infected with MOI 50 of *S.*Tm*^wt^*, and analyzed as in C. L = ladder. **E-F**: Quantification of C-D. **G**: BMMCs were pre-treated with TAK-242 and/or MK-2206 for 30-45min and infected with MOI 50 of *S.*Tm*^wt^*. After 4h, IL-6 secretion was measured. **H-I**: BMMCs from TLR4 KO mice and corresponding WT BMMCs were left uninfected, or infected with MOI 50 of *S.*Tm*^wt^* SL1344 or the indicated TTSS-mutants. After 4h, BMMCs harboring vacuolar *S.*Tm were quantified (**H**), and after 24h IL-6 secretion was measured (**I-J**). **J** depicts an enlargement of **I** with independent statistical analysis (two-way ANOVA and Sidak’s posthoc test for both). Experiments were performed 2-3 times and mean ± SEM of pooled biological replicates is shown. Data was statistically analyzed with ANOVA and Dunnet’s posthoc test, using *S*.Tm*^wt^*-infected cells for comparisons to all other groups (A-B, E-G). For H-I, TLR4 KO BMMCs were compared with corresponding WT BMMC groups by two-away ANOVA with Sidak’s posthoc test.

Translocated SopB has in other cell types been shown to trigger the PI3-kinase-Akt pathway ^48–50^ which can be a potent pro-inflammatory signal. In line with this, *S*.Tm*^wt^* elicited highly elevated levels of phosphorylated Akt (P-Akt) in BMMCs, in sharp contrast to *S*.Tm^Δ*sopB*^, *S*.Tm^Δ4^ and *S*.Tm^Δ*invG*^ strains that all fail to express or translocate SopB (**Figure 5C, E**). Elevated P-Akt levels were also observed in Cyto D-pretreated *S*.Tm*^wt^*-infected BMMCs (**Figure S6E**), again uncoupling this response from bacterial internalization. Both baseline P-Akt levels, and the elevated levels of P-Akt induced by *S*.Tm*^wt^* could be abrogated by the selective Akt inhibitor MK2206 (**Figure 5D, F**). Pretreatment with this inhibitor also notably reduced IL-6 secretion from *S*.Tm*^wt^*-infected BMMCs (**Figure S6F**). Since lower levels of IL-6 were still detectable under this condition (**Figure S6F**), we hypothesized that TLR4-dependent (downstream of LPS sensing) and Akt-dependent (downstream of the *S*.Tm TTSS-1 effectors) signals may combine to elicit cytokine secretion. The experimental data supported this notion; pretreatment with TAK242 + MK2206 caused more pronounced reduction of IL-6 secretion from *S*.Tm*^wt^*-infected BMMCs than either of the inhibitors alone (**Figure 5G**).

From these data, we postulated a two-step activation mechanism whereby extracellular *S*.Tm can be sensed by TLR4, generating a slow and weak MC cytokine response. Upon *S*.Tm invasion on the other hand, this TLR4 signal combines with signals elicited by the TTSS-1 effectors SopB/E/E2 (including SopB-triggered Akt-phosphorylation), leading to swift and full-blown MC cytokine secretion. To formally test this model, we established BMMCs from *Tlr4^−/−^* (C57BL/6J) mice. As expected, *S*.Tm invaded WT and *Tlr4^−/−^* BMMCs with similar proficiency and in a TTSS-1-dependent manner (**Figure 5H**). At 4h p.i., *S*.Tm*^wt^* and *S*.Tm single-effector mutants could still elicit IL-6 secretion also from *Tlr4^−/−^*BMMCs, but *S*.Tm^Δ*invG*^, *S*.Tm^Δ4^ and *S*.Tm^Δ*sopBEE*2^ did not (**Figure S6G**). When extending the time frame considerably (analysis at 24h p.i.), *Tlr4^−/−^* BMMCs were found to produce ∼2-3-fold lower levels of IL-6 than WT BMMCs in response to *S*.Tm*^wt^* (**Figure 5I**). Strikingly, the non-invasive *S*.Tm strains essentially failed to elicit a response altogether in the absence of TLR4 (**Figure 5I-J**). We conclude that MCs can use a two-step activation mechanism and tune their cytokine secretion output to differentiate between a non-invasive vs. invasive *S*.Tm encounter.

### The broad scale cytokine response from *Salmonella*-infected mast cells can promote myeloid cell survival and differentiation

To define the global MC gene expression changes elicited by invasive and non-invasive *S*.Tm, we next performed RNA Sequencing (RNASeq) of BMMCs left uninfected, exposed to *S*.Tm*^wt^*, or to *S*.Tm^Δ*invG*^ (MOI 50; analysis at 4h p.i.). Principal component analysis (PCA; based on all transcripts) illustrated a clear separation of the three sample groups along the PC1 axis (**Figure S7A**). Pairwise comparisons of either *S*.Tm*^wt^*-infected vs. uninfected MCs, or *S*.Tm*^wt^*- vs *S*.Tm^Δ*invG*^-infected MCs, showed that invasive infection brought about pronounced upregulation of a large panel of transcripts (**Figure S7A-F**; 3471 transcripts significantly upregulated between *S*.Tm*^wt^*-infected and uninfected samples). Among these were mRNAs encoding IL-6, TNF, and IL-13, hence validating our results by qPCR and ELISA (**Figure 2-4**). Additional cytokine/chemokine transcripts induced by invasive infection were e.g. *Ccl3*, *Ccl4*, *Ccl7*, *Cxcl10*, and *Csf2* (encoding GM-CSF) (**Figure 6A**). Another set of transcripts exemplified by *Il1b* showed equal levels of induction by *S*.Tm*^wt^* and *S*.Tm^Δ*invG*^ (**Figure 6A**), again corroborating the earlier qPCR results (**Figure S3E,K**).

**Figure 6.**
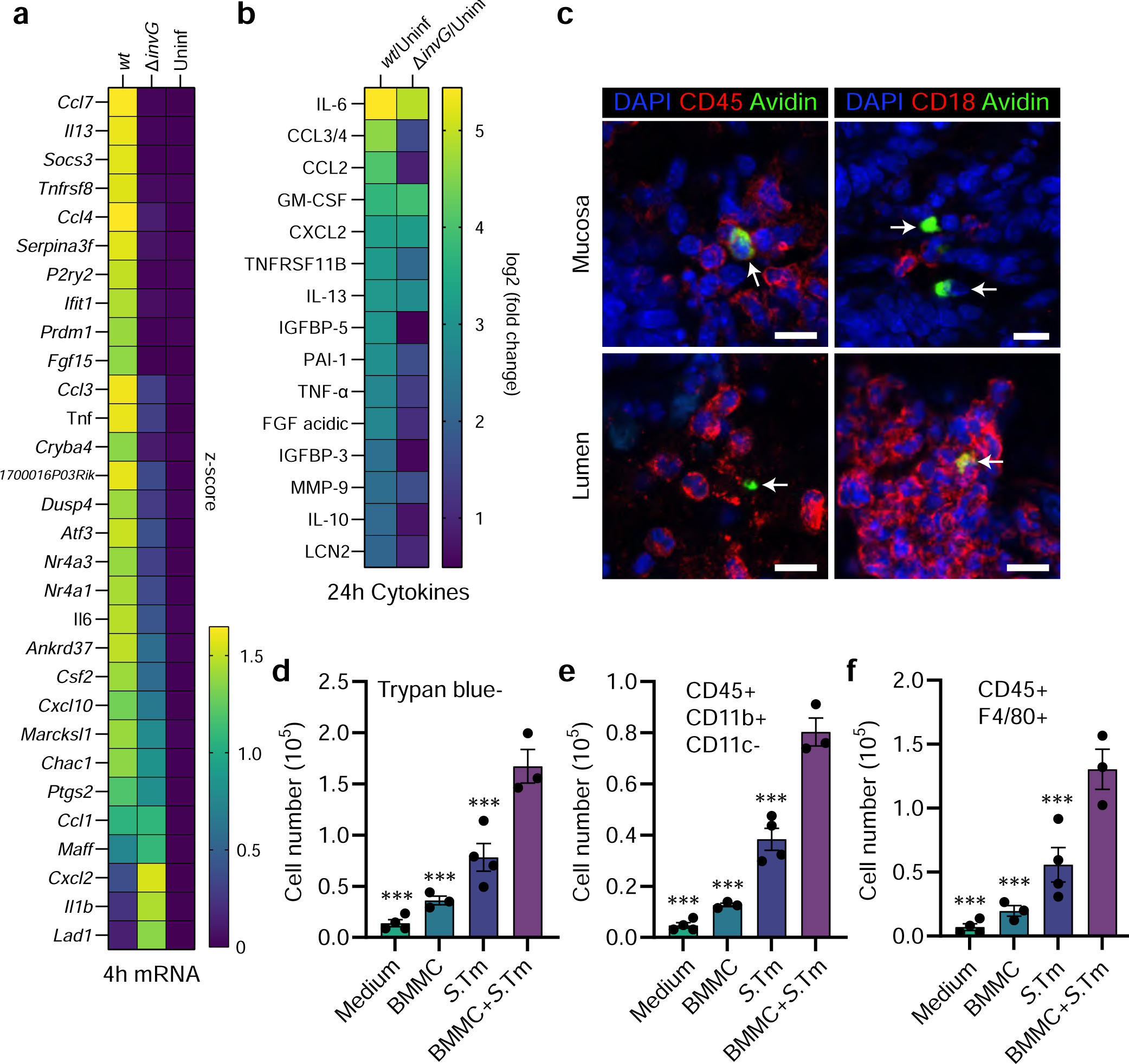
*Salmonella* induces a broad transcriptional and cytokine secretion response in mast cells with functional consequences on myeloid cells. **A**: RNA sequencing of BMMCs left uninfected, or infected with MOI 50 of *S.*Tm*^wt^*or *S*.Tm^Δ*invG*^ SL1344 for 4h, presented as the top 30 significantly upregulated genes between *S*.Tm*^wt^*-infected and uninfected control, displayed as a z-score-transformed heatmap. Genes are sorted by the formula “*S*.Tm*^wt^* - *S*.Tm^Δ*invG*^” to highlight differences between those two groups. **B**: Heatmap of relative log2 fold changes between the indicated groups, derived from a cytokine array of 24h supernatants from BMMCs infected with MOI 50 of *S.*Tm*^wt^* or *S*.Tm^Δ*invG*^ SL1344. **C**: Representative IF images of MCs in close contact to CD45+ and CD18+ immune cells in the *S.*Tm*^wt^* SL1344-infected intestinal mucosa and lumen at 48h p.i.. Arrows indicate MCs, scale bars: 10 µm. **D**: Trypan blue-based live cell counts of bone marrow nucleated cells, cultured for 7 days in base medium supplemented with either medium alone, 24h uninfected BMMC supernatant, *S.*Tm inoculum-conditioned supernatant, or supernatant of BMMCs infected with *S.*Tm*^wt^*for 24h. **E-F**: Similar setup as in C, but quantification of the total number of CD45^+^ CD11b^+^ CD11c^−^ cells (containing monocytes) (**E**) and CD45^+^ F4/80^+^ cells (containing macrophages) (**F**) in bone marrow cultures treated with the indicated supernatants, or base medium alone. For C-E, data was statistically analyzed with ANOVA and Dunnet’s posthoc test, using “BMMC+*S*.Tm” for comparisons to all other groups.

In a subsequent approach, a semi-quantitative array detecting 111 cytokines was used to screen supernatants of BMMCs either left uninfected, exposed to *S*.Tm*^wt^*, or to *S*.Tm^Δ*invG*^ (MOI 50; analysis at 24h p.i.) (**Figure 6B**). 15 cytokines showed a >4-fold increase between *S*.Tm*^wt^*-infected and uninfected BMMCs, while 5 were also >4-fold increased between *S*.Tm^Δ*invG*^-infected and uninfected samples (**Figure 6B**). The highest fold-change between *S*.Tm*^wt^*-infected and uninfected BMMCs was seen for IL-6, followed by CCL3, CCL2, GM-CSF, and then by IL-13 and TNF, in full agreement with our previous ELISA analyses (**Figure S2C-D, H-J**). Overall, this cytokine profile can be interpreted as a mixed pro-inflammatory (e.g. IL-6, TNF) and immunomodulatory (e.g. IL-10, PAI-1) output that may foster recruitment of granulocytes (e.g. CCL3, CXCL2), dendritic cells and monocytes (e.g. CCL2, CCL3), as well as promote survival and differentiation of macrophages (e.g. GM-CSF). Indeed, when further scrutinizing gut tissue harvested from *S*.Tm-infected mice, mucosal and luminal MCs were found to be surrounded by high densities of CD45 (broad blood cell marker) and CD18 (marker for monocytes, macrophages, and granulocytes) positive cells (**Figure 6C**).

In light of these observations, we finally examined the functional capacity of infected MC secretions, by culturing nucleated mouse bone marrow cells for 7 days in a base medium (see Materials and methods) that on its own failed to support cell survival (**Figure 6D**). The base medium was mixed with 24h conditioned supernatants from either uninfected BMMCs, the *S*.Tm*^wt^*inoculum alone, or *S*.Tm*^wt^*-infected BMMCs. Supernatants from naïve BMMCs had minimal impact on the bone marrow cells, but *S*.Tm-conditioned supernatants enhanced cell survival and macrophage differentiation to some extent (**Figure 6 D-F**; flow cytometry gating shown in **Figure S8**). However, both of these stimulatory effects were drastically higher for supernatants harvested from *S*.Tm*^wt^*-infected BMMCs. We conclude that cytokines and other soluble factors secreted by BMMCs upon *S.*Tm infection enhance survival of bone marrow-derived progenitors and may act in concert with soluble *S*.Tm components to promote myeloid cell differentiation.

## Discussion

This work establishes that MCs, common innate immune cells of mucosal tissues, can tune their cytokine response to extracellular vs. invasive forms of the prototype enterobacterium *S*.Tm and close relatives. Prior studies have shown that MCs respond to PAMPs of both gram-negative and -positive bacteria through TLRs ^6–9^. We corroborate these findings, showing that *E. coli* LPS, or non-invasive *S*.Tm strains, trigger a TLR4-dependent cytokine response. Importantly however, this weak response, observed also upon exposure to non-invasive *E. coli* and *Y. pseudotuberculosis* strains, vastly undershoots the maximal capacity of MCs to initiate *de novo* transcription and cytokine production. Instead, a full-blown MC response requires a second activation step directly linked to TTSS-1-dependent *S*.Tm invasion effectors. Through this wiring, MCs appear capable of informing their surrounding of a bacterium’s virulence potential. Experiments separating *S*.Tm-infected from neighboring MCs localized the vigorous MC response specifically to the bacterium-invaded cell population. This excludes that the second MC activation step is elicited by DAMP release from cells damaged by the infection (further supported by ^16^). Moreover, an *S*.Tm strain that retains TTSS-1 translocon function in the absence of TTSS-1 effectors (*S*.Tm^Δ4^) did not recapitulate the MC response to *S*.Tm*^wt^*. This also excludes that the second activation step comprises sensing of plasma membrane perturbation, as has been noted for MCs exposed to bacterial cytolysins ^14–16^. Rather, *S*.Tm single- and multiple gene mutant infections, and pharmacological inhibition assays, point to the translocated TTSS-1 effectors SopB, SopE, and SopE2 as responsible for the invasion-linked signal(s). These effectors have previously been linked to proinflammatory transcription in epithelial cells ^48^. SopB alters plasma membrane phosphatidylinositol phosphate (PIP) pools to generate e.g. PI(3,4)P_2_, through phosphatase and phosphotransferase activities ^51,52^. This recruits multiple kinases and promotes phosphorylation/activation of Akt, with consequences on host cell transcription and cell survival ^48–50,53^. We found that MCs exposed to *S*.Tm*^wt^* exhibited elevated levels of phospho-Akt, while this was not observed for strains incapable of SopB translocation (*S*.Tm^Δ*invG*^, *S*.Tm^Δ4^, *S*.Tm^Δ*sopB*^). MC pretreatment with the Akt inhibitor MK2206 also attenuated the MC cytokine response. Notably, however, the magnitude of the SopB contribution varied between MC models, whereas a *S*.Tm^Δ*sopBEE*2^ triple mutant consistently elicited a similar low response as the TTSS-1-deficient strains. SopE/E2 can, in partial redundancy with SopB, activate Rho GTPases such as Rac1 and Cdc42, which promote *S*.Tm invasion, but may also elicit proinflammatory transcription through MAP kinase and/or Nod1 pathways ^48,54,55^. It therefore seems most likely that invasive *S*.Tm spark a mixed transcription-stimulating signal in MCs through translocated SopB/E/E2, triggering both Akt activation, and additional pathways redundantly activated by the three effectors.

Our study also weighs in on the question if MCs permit bacterial internalization. Previous reports suggested that enterobacteria like *S*.Tm are not efficiently taken up by MCs, and thereby fuel a more restricted MC response than for example *Staphylococcus aureus* ^6,21^. However, we demonstrate here that *S*.Tm*^wt^* grown under TTSS-1-inducing conditions efficiently invade and establish an intracellular niche (evident by *ssaG*-GFP expression and TEM analyses) within both mouse (BMMC, PCMC) and human (LUVA) MC models. By contrast, genetically TTSS-1-deficient bacteria (*S*.Tm^Δ*invG*^), or *S*.Tm*^wt^*grown under non-TTSS-1-inducing conditions (overnight culture in LB with vigorous shaking) had an essentially non-invasive phenotype in cultured MCs. The discrepancy between our and previous findings are likely explained by how the *S*.Tm inoculum was prepared. In either case, our results favor the conclusions that i) MCs have a minimal inherent capacity for phagocytosis of enterobacteria, but that ii) they are highly susceptible to TTSS-1-mediated active *S*.Tm invasion.

According to the two-step activation mechanism proposed here, MCs are capable of a graded response to bacterial infection that depends on the pathogeńs invasive capacity. Notably, this response does not include overt degranulation. IgE-mediated activation of FcεRI-receptors elicits prompt MC degranulation ^56^, but this is typically not seen upon bacterial detection, although some reports have linked degranulation to certain microbial stimuli ^57,58^. Our data do not exclude that the *S*.Tm TTSS-1 effector SptP can have a MC degranulation-suppressing effect “in trans”, that is when degranulation is stimulated by other means, as has been proposed by others ^40^. However, neither *S*.Tm*^wt^*, nor *S*.Tm^Δ*sptP*^, elicited above-background degranulation of cultured MCs, suggesting that degranulation plays a minimal role in the immediate MC response to this bacterium. Instead, extracellular *S*.Tm, *E. coli*, or *Y. pseudotuberculosis*,or pure *E. coli* LPS sensed through TLR4, gave rise to a weak production of cytokines and chemokines. Upon two-step detection of invasive *S*.Tm through both TLR4- and SopBEE2-elicited pathways, this cytokine/chemokine response was both faster and more vigorous. While not the main focus of this study, we also detected type I interferons in response to *S*.Tm*^wt^*, which is in line with earlier proposals linking intracellular localization of a pathogen to boosted type I interferon production ^6^. Among the secreted proteins strongly stimulated by invasive *S*.Tm were both typical pro-inflammatory (e.g. IL-6, TNF) and immunomodulatory (e.g. IL-10) cytokines, chemokines linked to granulocyte (e.g. CCL3, CXCL2) and monocyte (e.g. CCL2, CCL3) recruitment, and growth factors (e.g. FGF, GM-CSF). Indeed, we could also substantiate that MCs in the *S*.Tm-infected gut frequently colocalize with a variety of other immune cell types, and that supernatants from *S*.Tm*^wt^*-infected MCs boosted immune cell progenitor survival, as well as myeloid cell differentiation.

How can this MC response be integrated into the current understanding of enterobacterial infection *in vivo*? In the mouse model experiments, MCs were found in the intestinal submucosa and mucosa prior to per-oral infection, and the mucosal MC population expanded further in the infected group. We also found evidence for MCs transmigrating into the *S*.Tm-filled lumen, coming in direct contact with dense bacterial populations in the process. It is well established that *S*.Tm express TTSS-1 in the lumen and use this to invade intestinal epithelial cells and elicit acute inflammation through epithelial inflammasome activation ^29,59^. During transition across the intestinal mucosa, TTSS-1 expression is gradually downregulated ^60^, generating a stealthier *S*.Tm phenotype within deeper tissues. Hence, it appears most plausible that a two-step sensing of TTSS-1-proficient *S*.Tm by MCs is predominantly relevant in the bacterium-laden gut mucosa and lumen, and that this generates a secretory MC output that helps shape the local microenvironment. The triggering of inflammation is a double-edged sword during enterobacterial infection. In the case of *S*.Tm, the acute host response to bacterial invasion can restrict pathogen translocation across the gut mucosa, but when dysregulated also disrupt the epithelial barrier and promote pathogen overgrowth ^61,62^. During intraperitoneal *S.*Tm infection, MC-derived TNF was in fact found to worsen disease outcome and fuel bacterial colonization ^63^. Hence, it seems critical for MCs (as well as other mucosal cell types) to carefully adjust their inflammation-modulatory output to the properties of the intruder and thereby foster a protective, rather than deleterious, counter response. This study provides proof-of-principle evidence that MCs can indeed grade their cytokine output by combining classical PAMP sensing with effector-triggered immunity ^64,65^, which enables them to differentiate between non-invasive and invasive enterobacterial infection.

## Materials and Methods

### Mice for mast cell culture

For experiments involving *Tlr4^−/−^* BMMCs, B6(Cg)-*Tlr4^tm^*^1^*^.2Karp^*/J (#029015) and corresponding C57BL/6J WT controls (#000664), 8 weeks old mice were purchased from The Jackson Laboratory. For all other experiments involving bone marrow or BMMCs, C57BL/6 WT mice (8-14 weeks old), bred and maintained at the National Veterinary Institute (SVA, Uppsala, Sweden) were used. The experimental procedures were approved by the local animal ethics committee (Uppsala djurförsöksetiska nämnd, Dnr 5.8.18-05357/2018). Whenever possible, remaining bones from WT C57BL/6 mice used as controls in other experiments were acquired for culturing BMMCs.

### Mouse infections

Eight-week-old female CBA mice (Charles River) were acclimatized to the new environment for one week before infection. 24h before infection, mice were pretreated with 20mg streptomycin per oral gavage. Mice were deprived of food and water for 4h prior to infection by oral gavage with 3.0-7.5×10^6^ CFU *S.*Tm in 100µl Dulbecco’s phosphate-buffered saline (PBS). For infection, *S.*Tm were grown in LB overnight, diluted 1:20 and re-grown for 4h before diluting bacteria in PBS for infection. The infection dose was confirmed by viable counts on streptomycin plates. Mice were monitored frequently for signs of unhealth. The experiments were approved by the local animal ethics committee (Umeå djurförsöksetiska nämnd, Dnr A27-17) and mice were housed in accordance with the Swedish National Board for Laboratory Animals guidelines.

### Cryosectioning of caecum, immunofluorescence, and toluidine staining of paraffin sections

For preservation in paraffin, tissues were fixed in 4% PFA for 6-12h at room temperature, rinsed in PBS and 70% ethanol, and immediately dehydrated and paraffinized in Tissue-Tek®VIP (SAKURA). Tissues were embedded with HistowaxTM paraffin (Histolab) with Tissue-Tek®TEC (SAKURA) and kept at 4°C until sectioning. For cryo-embedding, tissue pieces were placed in 4% PFA/ 4% sucrose/ PBS solution for 6-12h at room temperature followed by incubation in 20% sucrose/ PBS (overnight 4°C). Excess liquid was removed, tissue placed in OCT, flash frozen in liquid N_2_ and stored at −80°C. Cryosections of 20 µm were cut on a Cryostat CryoStar NX70 (Epredia) with at least 40µm distance between sections, placed on Superfrost Plus Adhesion Microscope Slides (Thermo Fisher Scientific, #J1800AMNT) and dried >16h. Sections were rehydrated in PBS (Gibco, #70013-016 or #14190144) for 5min and permeabilized for 3min in PBS/ 0.1% Triton TX-100. Slides were stained with 2.5µg/ml DAPI (Sigma-Aldrich, #D9542), 4U/ml phalloidin-A647 (Thermo Fisher Scientific, #A22287) and 10µg/ml avidin-A488 (Thermo Fisher Scientific, #A23170) for 40min. After washing 3x in PBS for 3min, mounting was done with Mowiol 4-88 (Sigma-Aldrich, #81381) and slides were dried overnight before storing at 4°C. MCs were counted as avidin+ cells per caecum section. For staining of *S*.Tm, slides were blocked in 10% normal goat serum (Sigma-Aldrich, #G9023) in PBS for 30min after the permeabilization step, followed by 40min incubation with *Salmonella* O Antiserum Factor 5, Group B in blocking buffer. After washing, slides were stained as described above but including Goat-α-rabbit-IgG(H+L)-Cy3 (Molecular probes, #A10520) 1:200 in PBS. For staining with CD18 and CD45, Goat-α-Rat-IgG(H+L)-AF647 (Invitrogen, #10666503) was used without phalloidin instead. For staining of *Salmonella*, all washing steps were performed only for 1min to avoid loss of bacteria. For toluidine stained tissue sections, 5µm paraffin tissue sections were cut in Microm HM360 (Zeiss), placed on SuperFrost slides and dried (50°C for one hour or overnight at 37°C). Sections were deparaffinized in xylene at 60°C (3 x 10min) and rehydrated through a graded series of alcohol. Sections were incubated with 0.1% toluidine in 1% NaCl pH 2.3-2.5, water, 95% ethanol, and mounted in dibutylphtalate xylene (DPX) (Sigma-Aldrich) after dehydration in 99% ethanol and xylene. All primary antibodies and their dilutions used in this study are shown in **S4 Table**.

### Bone marrow-derived mast cell culture

Bone marrow from tibiae and femurae of one mouse per culture was flushed out with PBS, washed 1x at 300 x g for 7min and filtered through a 70µm cell strainer. Cells were resuspended in 50ml BMMC culture medium consisting of 90% DMEM (Fisher Scientific, #31966047), 10% heat-inactivated FBS (Thermo Fisher Scientific, #D5671) and supplemented with Penicillin-Streptomycin (100U/ml, 100µg/ml, Sigma-Aldrich, #P0781) and 10ng/ml recombinant IL-3 (Peprotech, #213-13). Medium was changed every 3-4 days to a fresh flask in the first 4 weeks of culture, maintaining a cell density of 0.5 x 10^6^ cells/ml. Afterwards, medium was changed every 3-7 days. BMMCs were used during 4-10 weeks of culture. *Tlr4^−/−^* BMMCs were used up to 4 months of culture. Cell numbers were determined by trypan blue (Thermo Fisher Scientific, #15250-061) exclusion and quantified by an automated cell counter (CountessTMII FL, Life Technologies).

### Mouse bone marrow cell culture

Mouse bone marrow was extracted as described above until the cell strainer filtering step. Erythrocytes were lysed by resuspending the pellet in 3ml of RBC Lysis Buffer (eBioscience, #00-4333-57) and incubating for 3min on ice. Nucleated bone marrow cells were washed in PBS and resuspended to 10^6^ cells/ml in BMMC culture medium. 900µl conditioned medium was placed in a 12 well plate and 100 µl of bone marrow cells were added. After 7 days, cells were harvested with cell lifters (Corning, #3008) and stained for flow cytometry as described below.

### Peritoneal cell-derived mast cell culture

After euthanizing the mouse, the abdomen was washed with 70% ethanol. The abdominal cavity was carefully cut open and the abdomen was rinsed with 5ml ice cold PBS by injecting and gently shaking the mouse. Lavage from each mouse was collected individually to avoid possible contamination. After collection, the lavage was centrifuged for 8min, 300 x g and resuspended in PCMC culture medium (BMMC culture medium with 50µM 2-Mercaptoethanol (Sigma-Aldrich, #M6250), 20 ng/ml SCF (Peprotech, #250-03) and IL-3. On the next day, cells were observed under the microscope and cells from separate mice were pooled into one culture. Medium was changed every 3-4 days and the cell density was adjusted to 0.5 x 10^6^ cells/ml.

### LUVA cell culture

LUVA cells (^45^; Kerafast, #EG1701-FP) are derived from untransformed CD34+ enriched mononuclear hematopoietic progenitor cells, obtained from a blood donor and cultured in the presence of IL-3, IL-6 and SCF. The cells were maintained in complete StemPro™-34 SFM (Thermo Fisher Scientific, #10639011), supplemented with 2mM L-glutamine and Penicillin-Streptomycin and subcultured every 3-4 days.

### *Salmonella* strains, plasmids, and culture conditions

All strains and plasmids used in this study can be found in **S1 and S2 Table**, respectively. In **Figure 2 and S2**, wild-type and mutant strains of *Salmonella enterica* serovar Typhimurium ATCC 14028 background were included ^66^. All other strains used in this study were of a SL1344 background (SB300; streptomycin resistant) ^67^. The *S*.Tm*^ΔsptP^*mutant was generated via transfer of a previously described deletion ^66^ from a *S.*Tm 14028 strain (C1172) to the SL1344 background by P22 transduction. For infections, *S.*Tm cultures were grown overnight at 37°C for 12h in LB 0.3 M NaCl with appropriate antibiotics on a rotating wheel incubator to optimize aeration, followed by subculturing in the same medium without antibiotics at a 1:20 dilution for 4h at 37°C. For non-virulence inducing conditions, *S*.Tm was grown for 20h at 37°C in LB under 190rpm shaking. Prior to infection, 1ml was spun down for 4min at 12000 x g and reconstituted in co-cultivation medium (cell-specific standard culture medium without IL-3 and antibiotics). *E. coli* strains were grown overnight in LB at 37°C and subcultured in LB for 2h at 37°C prior infection. *Y. pseudotuberculosis* were grown overnight in LB at 26°C. Bacteria were subcultured for 1h at 26°C in LB, containing 50 mM of CaCl_2_, followed by 1h shifting to 37°C. When using different bacterial species, ODs were adjusted to equal bacterial concentrations, based on CFU numbers. After infection, inocula were diluted 1:10^6^ and 50µl were plated onto agar plates with antibiotics when appropriate to enumerate colony forming units (CFU).

### Infection of mast cells with *S*.Tm

For all infections, mast cells (BMMCs, PCMCs or LUVA) were washed twice in PBS and resuspended in the respective culture medium without antibiotics (co-cultivation medium). In experiments performed for ELISA or RT-qPCR analysis, if not indicated otherwise, 500µl of 1 x 10^6^ mast cells/ml were added to 24 well plates (Sarstedt) and infected with 25µl inoculum, resulting in a multiplicity of infection (MOI) of 50, for 30min in 37°C, 5% CO_2_. Afterwards, gentamicin was added to a final concentration of 90µg/ml, leaving the intracellular bacteria intact. After an additional 3.5h, wells were harvested in tubes, centrifuged for 5min at 400 x g and supernatants and pellets were frozen separately. For degranulation assays, the incubation after addition of gentamicin was reduced to 30min and phenol red-free DMEM (Gibco, #31053028) was used in the co-cultivation medium. To generate samples for immunoblots, 1-2ml of cells were infected in 12- or 6 well plates with identical concentrations of gentamicin and bacteria but only 30min incubation after addition of gentamicin and prior to freezing, the pellets were washed with PBS. For experiments involving flow cytometry, 180µl of 0.556 x 10^6^ mast cells/ml were added to 96 well round bottom plates (Thermo Fisher Scientific, #163320) and infected by adding 20µl of bacteria to the indicated MOIs. After 30min incubation as above, plates were gently centrifuged for 3min, 200 x g, supernatants were discarded and the plates vortexed gently before adding 200µl/well of co-cultivation medium, containing 100µg/ml gentamicin. Plates were incubated for further 3.5h, washed 1x in 1% BSA (Sigma-Aldrich # A9418) in PBS (200 x g, 3min) and fixed in 2% PFA (Sigma-Aldrich, #158127) in the dark for 20-30min. After 1x washing, cells were resuspended in 1% BSA in PBS and stored at 4°C until flow cytometry analysis of mCherry or GFP-positive MCs. MK-2206 (Selleck chemicals, #S1078), TAK-242 (Sigma-Aldrich, #614316) or Cyto D (Sigma-Aldrich #C8273) were added diluted in medium 30-45min prior to infection. *E. coli* LPS (Sigma-Aldrich, #L4516) and flagellin (Sigma-Aldrich #SRP8029) were added diluted in medium and used as indicated in the respective figure legend. Transwell plates were from Sigma-Aldrich (#CLS3470).

### Microscopy

For fluorescence microscopy, 400μl of fixed BMMCs (0.5 × 10^6^ cells/ml) in 1% BSA in PBS were added to 24-well glass bottom plates (High performance #1.5 cover glass, Cellvis, #P24-1.5P). Images were acquired on a custom-built microscope, based on a Nikon Eclipse Ti2 body, in differential interference contrast (DIC) and green fluorescence channels with a Prime 95B (Photometrics) camera through a 100X/1.45 NA Plan Apochromatic objective (Nikon), using a X-light V2 L-FOV spinning disk (Crest Optics, Italy) and a Spectra-X light engine (Lumencor). Micro Manager was used for controlling the microscope (μManager plugin ^68^). For TEM, BMMCs infected with *S.*Tm MOI 50 for 4h (gentamicin present after 30min), or left uninfected, were fixed in 2.5% Glutaraldehyde (Ted Pella) + 1 % Paraformaldehyde (Merck) in PIPES pH 7.4 and stored at 4°C until further processed. Samples were rinsed with 0.1M PB for 10min prior to 1h incubation in 1% osmium tetroxide (TAAB) in 0.1M PB. After rinsing in 0.1M PB, samples were dehydrated using increasing concentrations of ethanol (50 %, 70 %, 95 % and 99.9 %) for 10min at each step, followed by 5min incubation in propylene oxide (TAAB). The samples were then placed in a mixture of Epon Resin (Ted Pella) and propylene oxide (1:1) for 1h, followed by 100% resin and left overnight. Subsequently, samples were embedded in capsules in newly prepared Epon resin and left for 1-2h and then polymerized at 60°C for 48h. Ultrathin sections (60-70nm) were cut in an EM UC7 Ultramicrotome (Leica) and placed on a grid. The sections were subsequently contrasted with 5% uranyl acetate and Reynold’s lead citrate and visualized with Tecnai™ G2 Spirit BioTwin transmission electron microscope (Thermo Fisher/FEI) at 80kV with an ORIUS SC200 CCD camera and Gatan Digital Micrograph software (both from Gatan Inc.). Cecal tissue cryosections stained for fluorescence microscopy were imaged using a LSM700 (Zeiss) confocal microscope, equipped with a Plan-Apochromat 40x/0.95 Korr M27 objective with pinhole set to 1 AU for each wave length, at the BioVis platform of Uppsala University. Fiji ^69^ was used for image analysis and distance measurements.

### Supernatant analyses by ELISA and β-hexosaminidase assays

BMMC and PCMC supernatants were analyzed by ELISA for IL-6, TNF and IL-13 (Invitrogen, # 88-7064-76, #88-7324-88, #88-7137-88). LUVA supernatants were assayed for IL-6 and TNF (Invitrogen, #88-7066-88, #88-7346-88). β-hexosaminidase was measured in fresh supernatants by transferring 20µl sample/well to a 96 well plate (Sarstedt) and adding 80µl of 1mM 4-Nitrophenyl N-acetyl-β-D-glucosaminide (Sigma-Aldrich, #N9376) in citrate buffer with pH 4.5. After 1h incubation at 37°C, the reaction was stopped with 200µl Na_2_CO_3_ buffer at pH 10 and absorption was measured at 405nm. Total β-hexosaminidase content was acquired by lysis with 1% Triton X-100 (Sigma-Aldrich, T8787) prior to the assay and for the respective infection experiment. MCs treated with 2mM calcium ionophore A23187 (Sigma-Aldrich, #C7522), were included as positive control for β-hexosaminidase release.

### Real-time quantitative PCR

Total RNA was isolated from frozen pellets and caecum tissue using the NucleoSpin RNA extraction kit (Machery-Nagel, #740955.5) or the RNeasy Plus Mini Kit (Qiagen, #74134). Equal concentrations of RNA (measured by NanoDrop and 1 µg for the caecum tissue) were reverse transcribed to cDNA using the iScript™ cDNA Synthesis Kit (Bio-Rad, #1708891) and stored at −20°C. cDNA was diluted 1:5 (for caecum 1:1 or 1:10) and qPCR was performed on a CFX384 Touch™ (Bio-Rad) with iTaq Universal SYBR (Bio-Rad, #1725121) using 1 µl of cDNA and 200 nM of reverse and forward primers (sequences are listed in **S3 Table**). PCR was performed according to the manufacturer’s instruction. If not indicated otherwise, data is shown as fold change (2^−ΔΔCq^) to uninfected MCs, normalized to *Hprt* transcription. Statistics were calculated based on ΔCq values.

### Flow cytometry

For detection of TLR4, 1 x 10^6^ BMMCs were resuspended in 2% BSA (Sigma-Aldrich) in PBS, containing 0.5µg/ml of Fc-block anti-CD16/32 (BD Life Sciences, #553142) and incubated for 5min at RT. The cells (100µl each) were either left unstained, or stained with 5µg/ml anti-CD284 (TLR4) antibody (clone UT41) conjugated with Alexa Fluor 488 (Invitrogen, #53-9041-82) or identical amount of mouse IgG1 kappa isotype control (Invitrogen, #MG120). After 30min incubation on ice, the cells were washed and resuspended in 2% BSA in PBS. For LPS staining, fixed cells were resuspended in 100µl of 0.1% saponin (VWR, Calbiochem, #558255), 1% BSA and 1:250 of *Salmonella* anti-serum in PBS and incubated for 1h at RT in the dark. After washing in 0.1% saponin, 1% BSA in PBS, the cells were incubated for 30min with 1:200 Goat-α-rabbit-IgG(H+L)-Cy3 in the dark and washed in 0.1% saponin again. Flow cytometry was performed using a MACSQuant VYB (Miltenyibiotec). For bone marrow analysis, cells were collected in 2% FBS in PBS containing fluorophore-conjugated antibodies targeting the following surface markers: CD45, CD11b, CD11c, F4/80. After 30min incubation, the cells were washed twice with 2% FBS in PBS, and resuspended in the same buffer for analysis. Flow cytometry was performed on a Cytoflex LX (Beckman Coulter). Data analysis was performed using FlowJo (BD Biosciences) version 10.8.1. Detailed information of all primary antibodies can be found in **S4 Table**.

### Cell sorting

After infection of BMMCs as described above for 4h, cells were washed in 1% BSA in PBS and resuspended in cold MACSQuant® Tyto® Running Buffer (Miltenyi Biotech, #130-107-207) to a concentration of 0.5 x 10^6^ cells/ml. All further steps were performed at 4°C. Cells were sorted on *ssaG*-GFP-expression in a MACS® GMP Tyto® Cartridge (Miltenyi Biotech #170-076-011) until purity of GFP-cells reached 99%. This procedure isolated GFP-BMMCs and enriched for GFP+ cells. After sorting, cells were washed in 1% BSA in PBS, pelleted and frozen until RNA extraction.

### Protein extraction and immunoblotting

Frozen pellets were resuspended in cold lysis buffer consisting of RIPA buffer (ThermoScientific, #89900) with protease inhibitors (Roche, #4693132001) and PhosSTOP™ (Roche, #4906845001), using 75µl per 1 x 10^6^ cells. After 30min incubation on ice, samples were centrifuged for 20min at 4°C max. speed to remove debris. Supernatants were transferred in fresh tubes, measured with the bicinchoninic acid assay (Pierce, #23227) and frozen until further use. Equal concentrations of protein (10-25µg) were mixed with 4x Laemmli buffer (Bio-Rad, #1610747) containing 5% 2-mercaptoethanol, boiled for 5min at 95 °C and loaded together with 5µl of ladder (Bio-Rad, #1610373) on 4–20% Mini-PROTEAN® TGX Stain-Free™ Protein Gels (Bio-Rad, #4568094 or #4568096). Gels were run for 25min at 200V and transferred to nitrocellulose (Bio-Rad, #1704158) with a Trans-Blot Turbo Transfer System (Bio-Rad), using the program for intermediate molecular weight. 1min activation of the gel and imaging of total protein on the blot was performed by a Gel Doc EZ Imager (Bio-Rad). Blots were blocked in Intercept blocking buffer (PBS, Li-cor, #927-70001) for 1h at RT, incubated 2h on RT or overnight at 4°C with antibodies for Akt (1:1000, Cell signaling, #9272) or P-Akt (Phospho-Akt (Ser473) (D9E) XP® Rabbit mAb (1:2000, Cell signaling, #4060) diluted in blocking buffer. After 3x washing in PBS-Tween (0.05%), blots were incubated with Goat-α-rabbit-IgG-HRP (1:10,000, Cell Signaling, #7074) for 1h at RT. After washing, blots were incubated with ECL Prime (Cytiva, #GERPN2232) for 5min at RT under shaking and imaged with a ChemiDoc MP (Bio-Rad). Two blots were run in parallel for Akt and P-Akt, antibody signal was normalized to total protein on the membrane respectively, and data was displayed as P-Akt of total Akt, normalized to untreated cells.

### RNA sequencing

Sequencing libraries were prepared from 500ng total RNA using the TruSeq stranded mRNA library preparation kit (Illumina Inc., #20020595,) including polyA selection. Unique dual indexes (Illumina Inc., 20022371) were used. The library preparation was performed according to the manufacturers’ protocol (#1000000040498). The quality of the libraries was evaluated using the Fragment Analyzer from Advanced Analytical Technologies, Inc. using the DNF-910 dsDNA kit. The adapter-ligated fragments were quantified by qPCR using the Library quantification kit for Illumina (KAPA Biosystems) on a CFX384 Touch instrument (Bio-Rad) prior to cluster generation and sequencing. Library preparation and sequencing was performed by the SNP&SEQ Technology Platform, a national unit within the National Genomics Infrastructure (NGI), hosted by Science for Life Laboratory, in Uppsala, Sweden (scilifelab.se/units/ngiuppsala). Sequencing was carried out on an Illumina NovaSeq 6000 instrument (NSCS v 1.7.5/ RTA v 3.4.4) according to the manufacturer’s instructions. Demultiplexing and conversion to FASTQ format was performed using the bcl2fastq2 (2.20.0.422) software, provided by Illumina. Additional statistics on sequencing quality were compiled with an in-house script from the FASTQ-files, RTA and BCL2FASTQ2 output files. The RNA-seq data were analyzed using the best practice pipeline nf-core/rnaseq. Detailed information about the analysis pipeline can be found here: ngisweden.scilifelab.se/bioinformatics/rna-seq-analysis and nf-co.re/sarek. For differential expression analysis, DESeq2 1.38.3 in combination with R 4.2.2 and RStudio 2022.12.0+353 were used. Transcriptome data can be accessed at Gene Expression Omnibus under GSE223601.

### Cytokine array

Proteome Profiler Mouse XL Cytokine Array (Bio-Techne, #ARY028) was used, following the manufacturer’s instruction for fluorescent detection with IRDye® 800CW Streptavidin (LI-COR, #926-32230) on an Odyssey CLx Infrared Imager. Raw images were cropped, converted to 16 bit with ImageJ and dot intensities were analyzed with Image Lab 6.1 (Bio-RAD). Ratios of log2 fold changes between groups were calculated.

### Statistical analysis

If not indicated otherwise, all graphs were plotted with Prism 9.5.1 (GraphPad) and statistical analysis was performed either with one-way analysis of variance (ANOVA) and Sidak’s posthoc test or two-factor ANOVA with Dunnett’s posthoc test in order to compare groups to either control or WT-infected MCs as indicated in the figure legends. Whenever appropriate, paired or unpaired t-tests or for data using mice, the Mann-Whitney U test were used. Significance levels were: * p < 0.05, ** p < 0.01 and *** p < 0.001. If not indicated otherwise, for every n, the mean of all MC wells infected with an individual subculture derived from an individual overnight culture serves as a single datapoint. If not indicated otherwise, every experiment was performed at least twice on different days.

## Acknowledgements

We thank members of the Sellin and Pejler laboratories for helpful discussions. We are grateful for support regarding TEM and fluorescence microscopy from BioVis, Uppsala University. This work was supported by the Knut and Alice Wallenberg Foundation (KAW 2016.0063 to MF and MES), the Swedish Research Council (2018-02223 to MES; 2016-00803 to JH), a Lennart Philipson Award (MOLPS, 2018, to MES), and the SciLifeLab Fellows program.

## Contributions

Conceptualization: CvB, GP, MES; Methodology: CvB, AF, PG, MLDM, EM-E, JE; Investigation: CvB, AF, OL, GIP, EM-E; Formal analysis: CvB, AF, EM-E; Interpretation: CvB, AF, PG, MLDM, OL, EM-E, JH, MF, GP, MES; Resources: JH, MF, GP, MES; Supervision: JH, MF, GP, MES; Funding acquisition: JH, MF, GP, MES; Visualization: CvB, MLDM; Writing - original Draft: CvB, MES; Writing - reviewing & editing: all authors.

## Data availability statement

The RNA seq data generated in this study have been deposited in the GEO under accession number GSE223601. The rest of the data are available in the article, Supplementary Information, or Source Data file. Source data are provided with this paper.

---

**S1 Figure.**
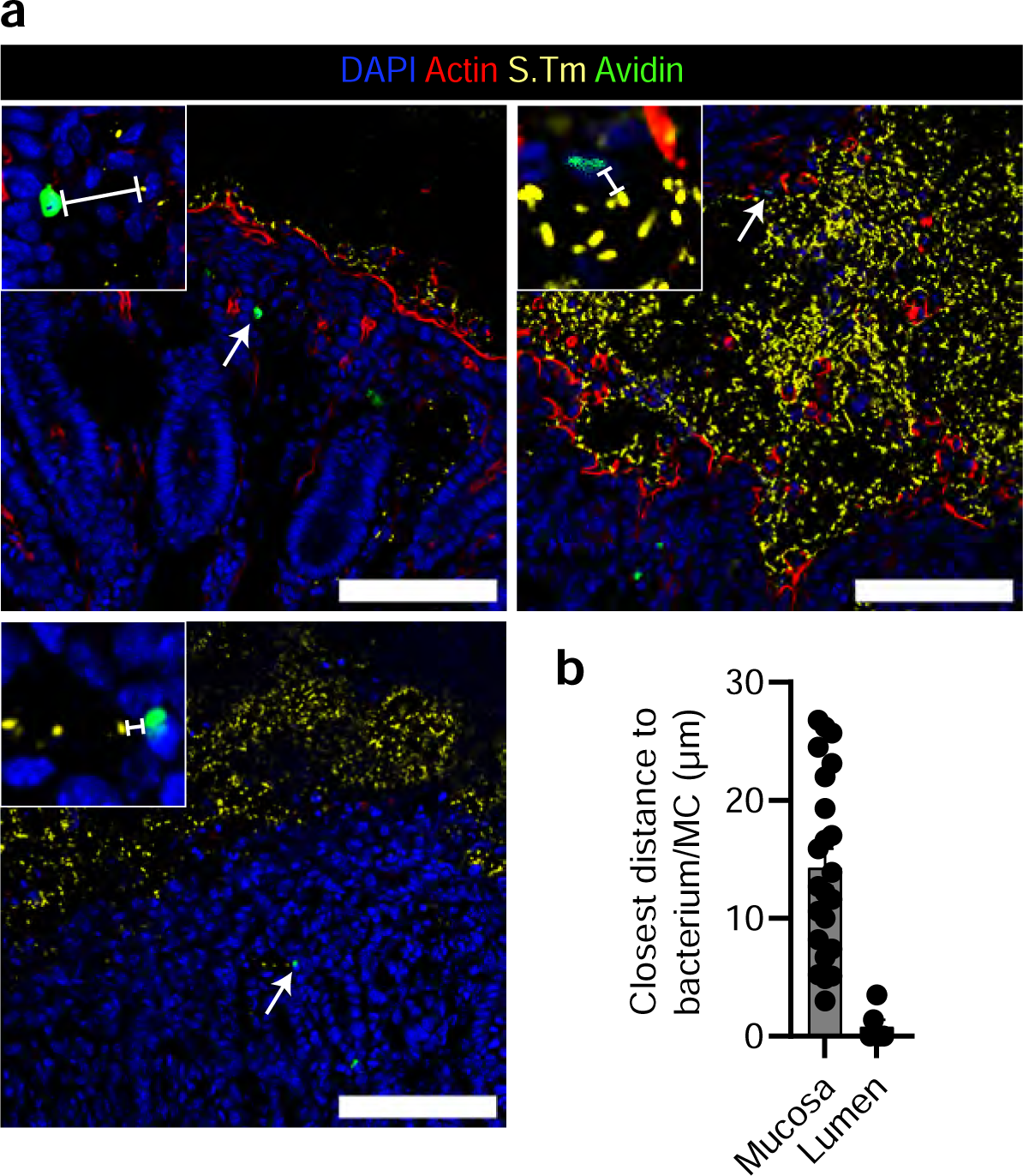
Mast cells can be found in close proximity to *S.*Tm. **A**: Representative IF images used in quantification of the distances of MCs to their closest bacterium. Scale bars are 100µm. Magnifications are in 50×50µm for the top left image and 25×25µm for the other two images. Arrows indicate magnified MCs. **B**: Quantification of distances to closes bacteria for individual MCs within fields of view containing both MCs and *S*.Tm. For individual MCs (23 in mucosa and 6 in the lumen), a straight line was drawn to the closes bacterium and the distance measured. Bars show mean ± SEM.

**S2 Figure.**
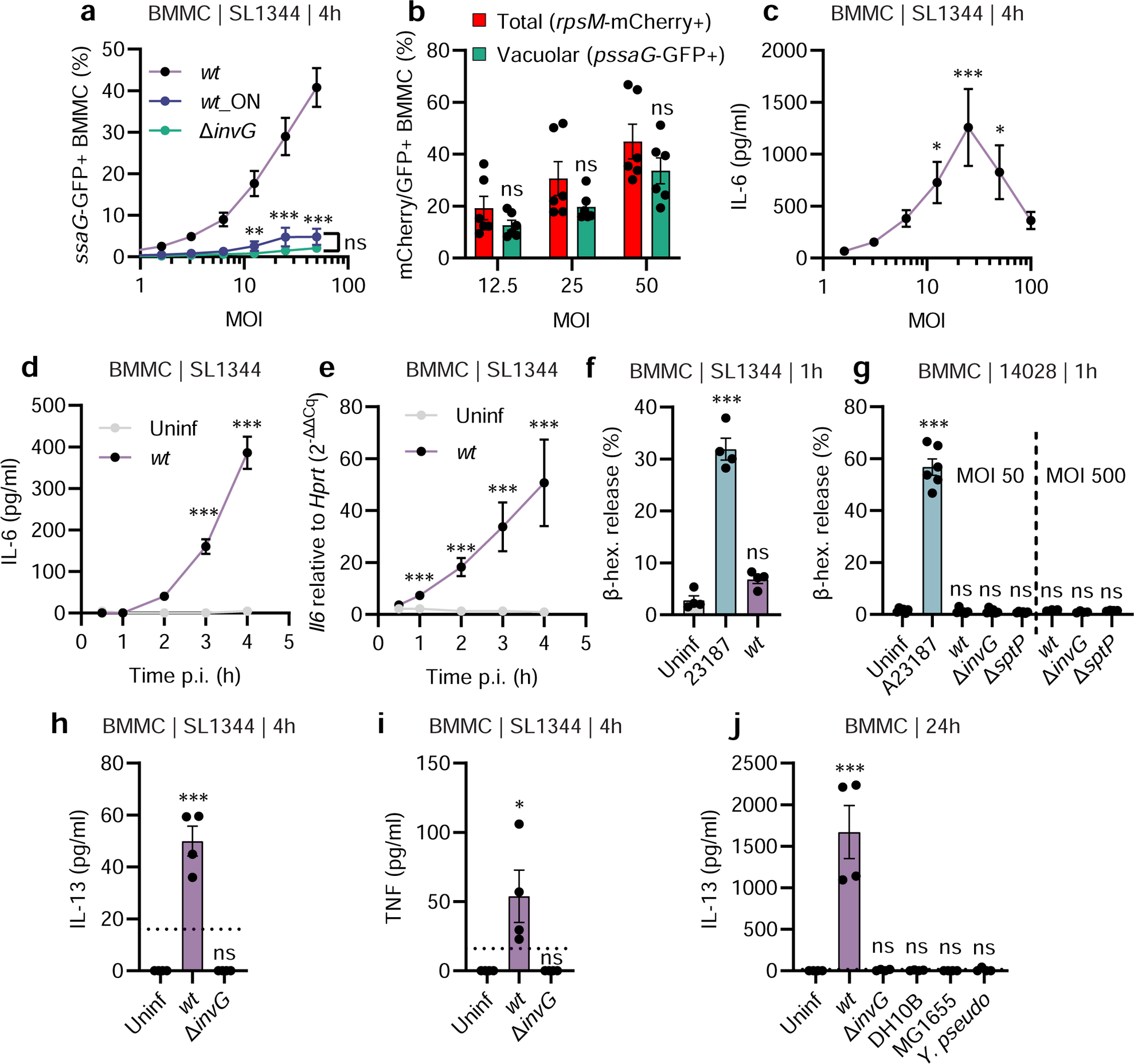
Mast cells respond to invasive *Salmonella* infection by cytokine gene transcription and secretion, but negligible degranulation, within the first hours. **A**: MOI-dependent quantification of BMMCs, harboring vacuolar *S.*Tm. “ON” indicates that the *S.*Tm inoculum is grown as an over-night stationary phase culture **B**: Quantification of BMMCs bound to or invaded by *S.*Tm (red bars) compared to BMMCs harboring vacuolar *S*.Tm (green bars).. **C**: MOI-dependent quantification of IL-6 secretion by BMMCs, infected with the indicated *S*.Tm SL1344 strains for 4h. **D-E**: IL-6 secretion **(D**) and *Il6* transcript levels (**E**) of BMMCs left uninfected or infected with MOI 50 of *S.*Tm*^wt^* SL1344 for the indicated time frames. **F**: β-hexosaminidase release from BMMCs as an indicator of degranulation 1h after infection with MOI 50 of *S.*Tm*^wt^* SL1344. A23187 served as positive control. **G**: Similar setup as in E, but with *S*.Tm*^wt^*or the indicated TTSS-mutants of strain 14028, at MOI 50 and MOI 500. **H-I**: IL-13 (**H**) and TNF (**I**) secretion from BMMCs, 4h after infection with MOI 50 of S.Tm*^wt^* SL1344 or the indicated TTSS-mutants. **J**: Secreted IL-13 of BMMCs infected with MOI 50 of S.Tm*^wt^* and *S*.Tm^Δ*invG*^ SL1344 as well as *E. coli* DH10B, *E. coli* MG1655 and *Y. pseudotuberculosis* for 24h. Every experiment was performed 2-3 times and mean ± SEM of pooled biological replicates is shown. Groups in A were statistically analyzed with two-way ANOVA and Tukey’s posthoc test (every group for each MOI to “*wt*” group). For C, F-I; one-way ANOVA together with Dunnet’s posthoc test was used with uninfected cells (in case of A “MOI 0”) were used for statistical comparison to all other groups. For D-E, two-way ANOVA with Sidak’s posthoc test was used to compare uninfected cells with *S*.Tm*^wt^*-infected cells within each time point. For ATCC 140828 infections, a Δ*malX* strain was used as *wt*.

**S3 Figure.**
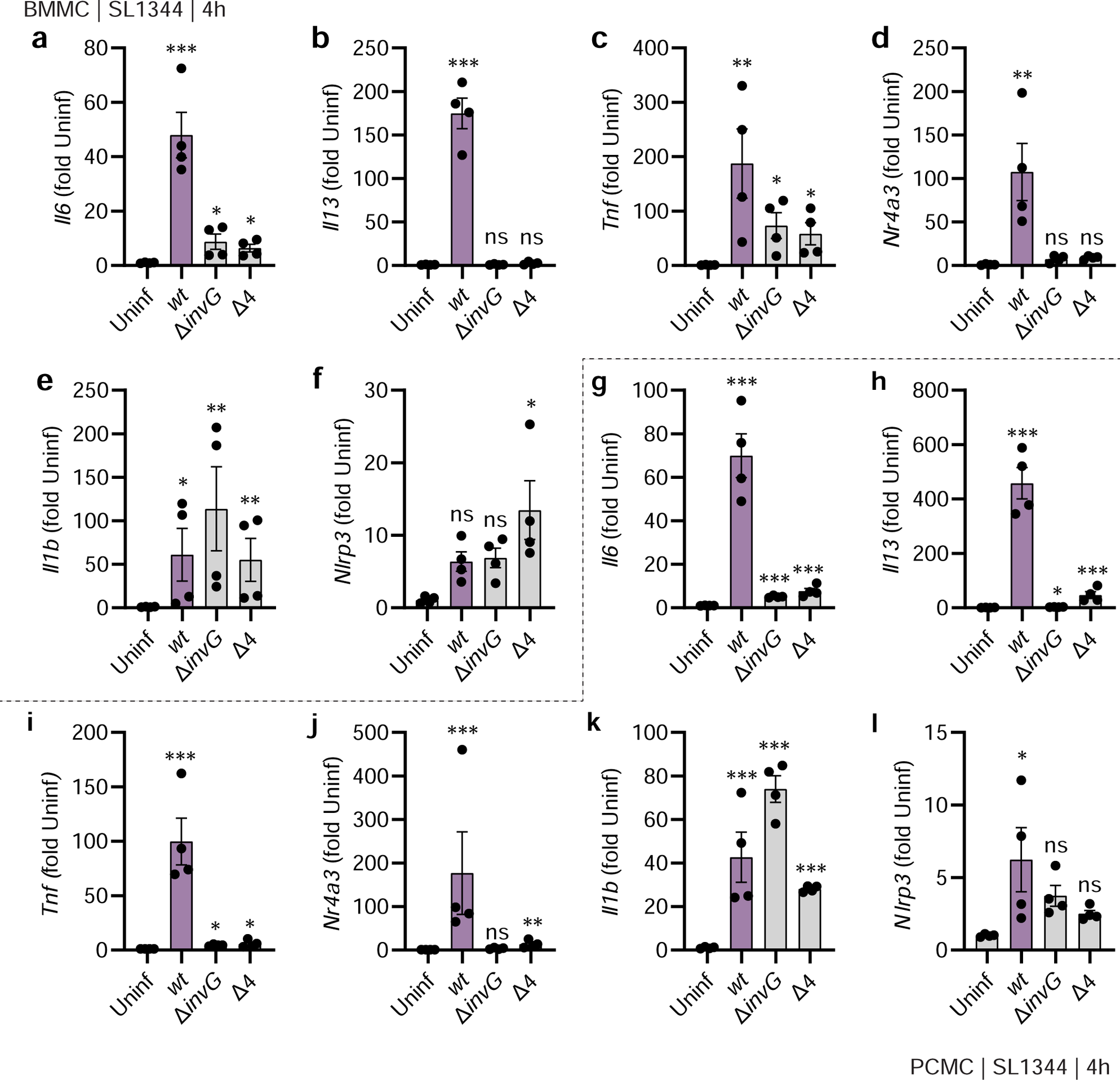
Distinct transcriptional profiles of mast cells infected with invasive vs. non-invasive *Salmonella*. **A-F** RT-qPCR quantification of transcript levels in BMMCs, 4h after infection with MOI 50 of *S.*Tm*^wt^* SL1344 or the indicated TTSS-mutants. **G-I:** Similar as in A-F, but PCMCs were used. Every experiment was performed 2-3 times and mean ± SEM of pooled replicates is shown. Data was statistically analyzed with ANOVA and Dunnet’s posthoc test, using uninfected infected cells for comparisons to all other groups. A selection of these data (A-D, G-J) are also summarized in heatmaps in Figure 2H-I.

**S4 Figure.**
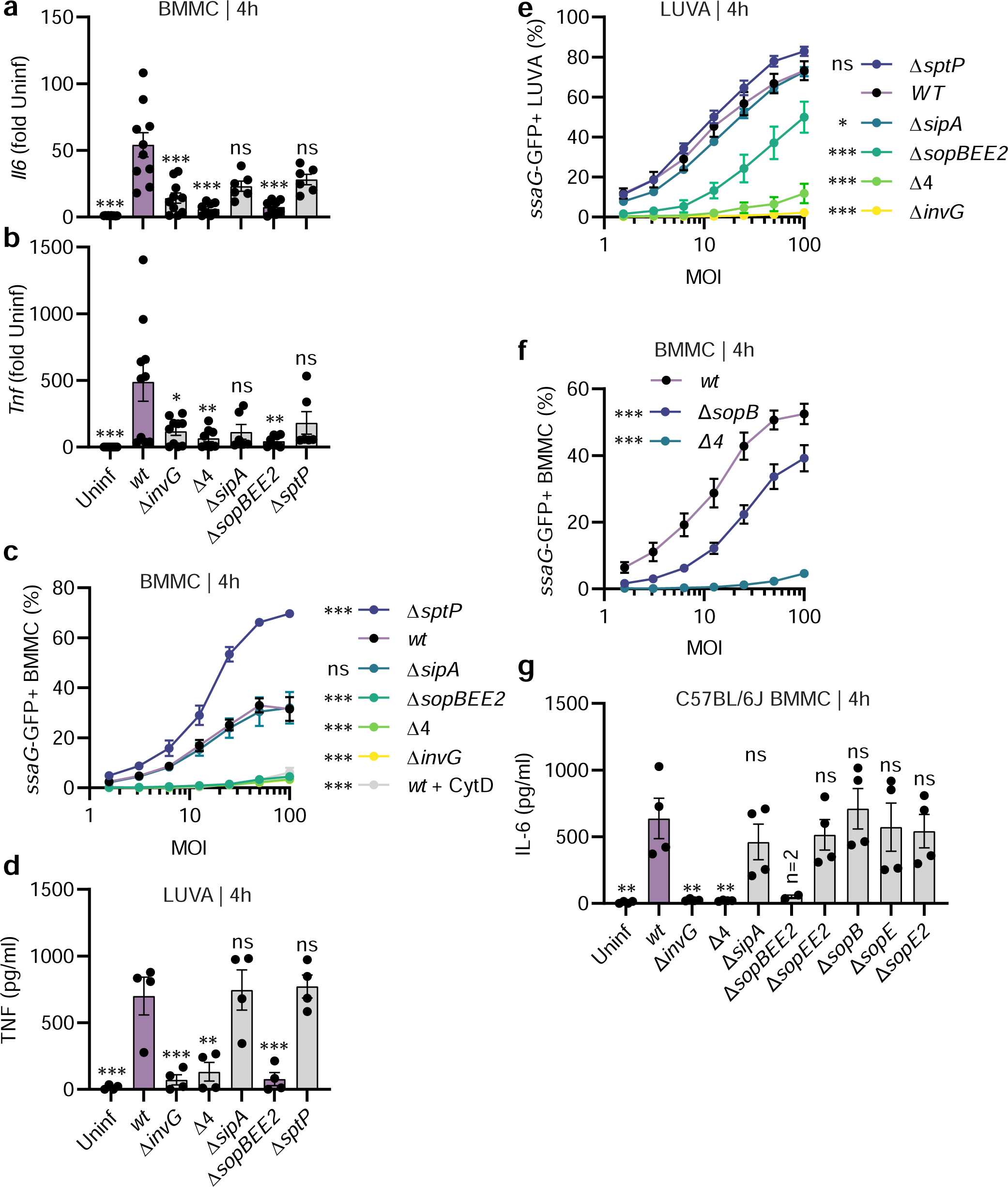
The TTSS-1 effectors SopB, SopE, and SopE2 promote murine and human mast cell cytokine secretion upon *Salmonella* infection. **A-B**: RT-qPCR quantification of *Il6* (**A**) and *Tnf* (**B**) transcript levels in BMMCs infected with MOI 50 of *S*.Tm*^wt^* SL1344 or the indicated TTSS-mutants for 4h. **C, F**: Quantification of the MOI-dependent frequency of BMMCs harboring vacuolar bacteria, 4h after infection with MOI 50 of S.Tm*^wt^* SL1344 or the indicated TTSS-mutants. For actin inhibition, BMMCs were pretreated with 1µM of Cyto D for 1h. **D**: TNF secretion from LUVA cells, infected with MOI 50 of S.Tm*^wt^* SL1344 or the indicated TTSS-mutants for 4h. **E**: Quantification of the MOI-dependent frequency of LUVA cells harboring vacuolar bacteria, 4h after infection with MOI 50 of S.Tm*^wt^* SL1344 or the indicated TTSS-mutants **G**: IL-6 secretion from C57BL/6J (Jackson) BMMCs infected with MOI 50 of *S*.Tm*^wt^* SL1344 or the indicated TTSS-mutants for 4h. Every experiment was performed 2-3 times and mean ± SEM of pooled biological replicates is shown. For A, B, D and G, data was statistically analyzed with ANOVA and the Dunnet’s posthoc test, using *S*.Tm*^wt^*-infected cells for comparison to all other groups. For C, E, and F, two-way ANOVA was used with Sidak’s posthoc test to compare the respective mutants and *S*.Tm*^wt^*.

**S5 Figure.**
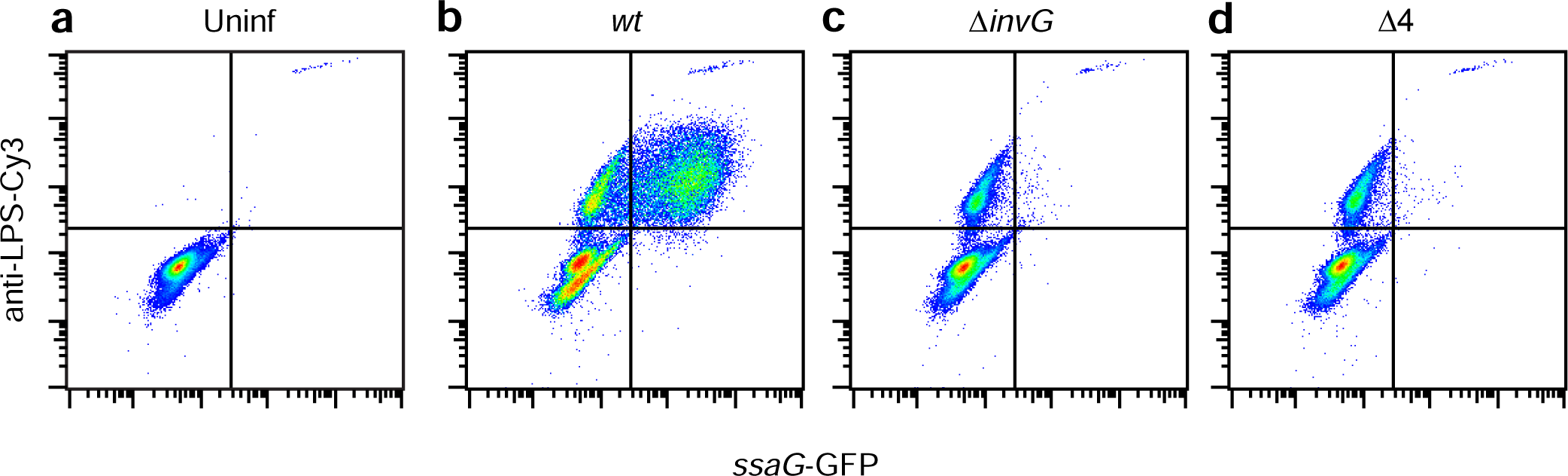
Flow cytometry gating for data shown in Figure 4B. **A**-**D**: Representative flow cytometry gating for quantification of MCs positive for vacuolar *S.*Tm (*ssaG*-GFP+) and/or *S*.Tm LPS. BMMCs were infected with MOI 50 of *S*.Tm*^wt^* SL1344 or the indicated TTSS-mutants for 4h prior to analysis by flow cytometry.

**S6 Figure.**
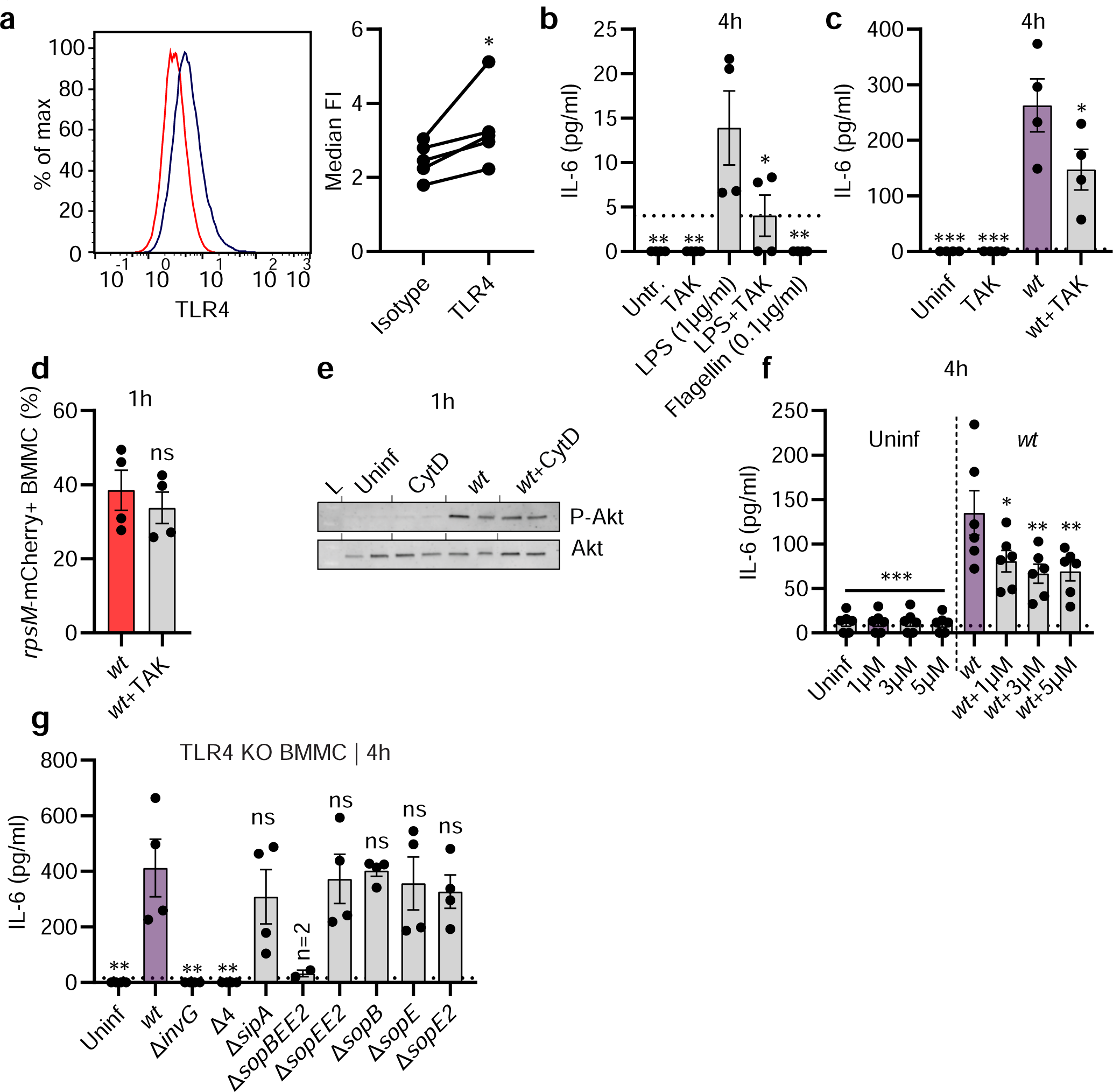
Mast cells express surface TLR4 and react to *Salmonella* through pathways involving TLR4 and Akt. **A**: Representative surface detection of TLR4 (blue) and isotype control (red) in BMMCs. Pairwise comparisons shown in right panel. **B**: IL-6 secretion from BMMCs 4h after treatment with LPS or recombinant flagellin and/or pretreatment for 30-45min with TAK-242. **C**: IL-6 secretion from BMMCs 4h after infection with MOI 50 of *S*.Tm*^wt^*SL1344 and/or pretreatment with TAK-242. **D**: Quantification of BMMCs harboring intracellular *S.*Tm 30min after infection. The indicated group was pretreated with TAK-242. **E**: P-Akt and Akt immunoblots of BMMCs after 1h infection with MOI 50 of *S*.Tm*^wt^* SL1344 and/or pretreatment with 200nM Cyto D. L = ladder. **F**: IL-6 secretion from BMMCs after 4h infection with MOI 50 of *S*.Tm*^wt^*SL1344 and/or pretreatment with MK-2206 for 30-45min. **G**: IL-6 secretion from *Tlr4^−/−^*BMMCs infected with MOI 50 of S.Tm*^wt^* SL1344 or the indicated TTSS-mutants for 4h. Every experiment was performed 2-5 times and mean ± SEM of pooled biological replicates is shown. E shows the results from 1 immunoblot. For A (paired) and D (unpaired) t-tests were used. For B, C, F and G, data was statistically analyzed with ANOVA and the Dunnet’s posthoc test, using *S*.Tm*^wt^*-infected cells (or LPS in case of B) for comparison to all other groups.

**S7 Figure.**
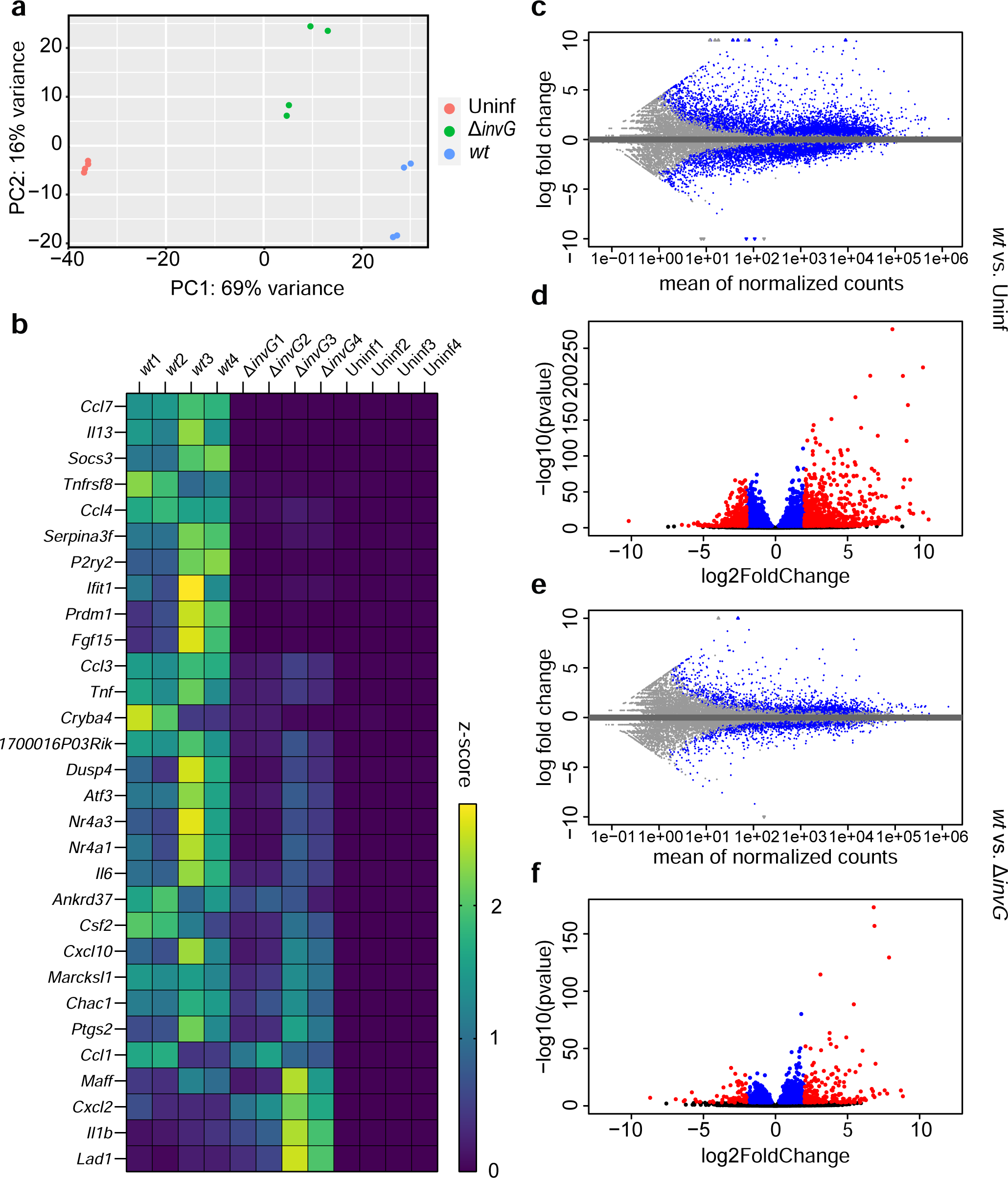
Mast cells react to invasive vs. non-invasive *Salmonella* with distinct transcriptional responses. **A**: Principal component analysis plot of the RNA sequencing data described in Figure 6A. **B**: Complete heatmap corresponding to Figure 6A, but showing z-scores for all four biological replicates in each group. Replicates “1” and “2” stem from one experiment, while “3” and “4” stem from a separate experiment, performed on a different day. **C**-**F**: Bland–Altman (**C**, **E**) and volcano (**D**, **F**) plots for comparisons between *S*.Tm*^wt^* and uninfected cells (**C**, **D**), or between *S*.Tm*^wt^* and *S*.Tm^Δ*invG*^ (**E**, **F**).

**S8 Figure.**
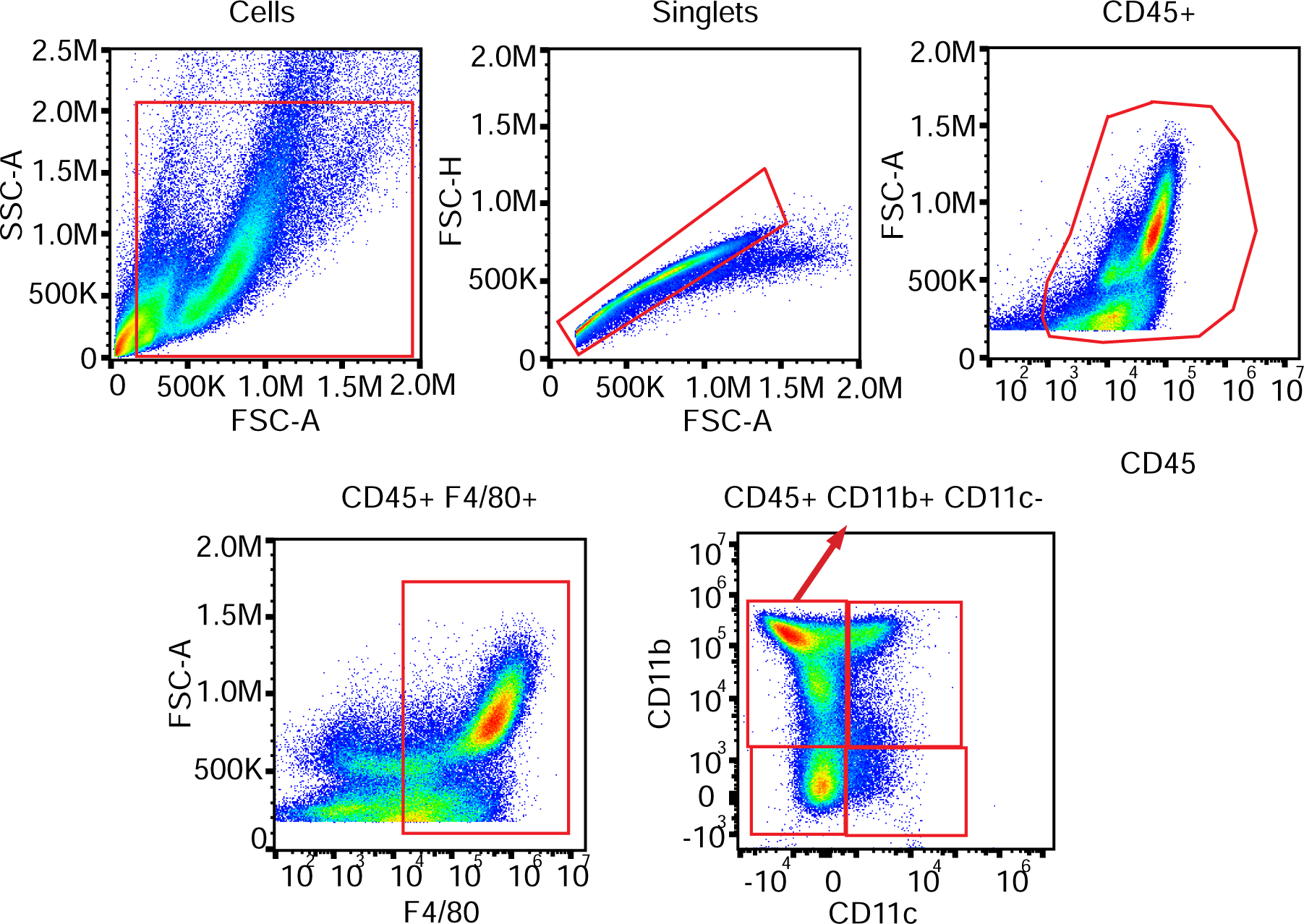
Flow cytometry gating for data shown in Figure 6D-E. Representative flow cytometry gating strategy, depicting one of the bone marrow nucleated cell samples, cultured for 7 days in base medium supplemented with supernatant from BMMCs infected with *S.*Tm*^wt^* for 24h.

**S1 Table.**
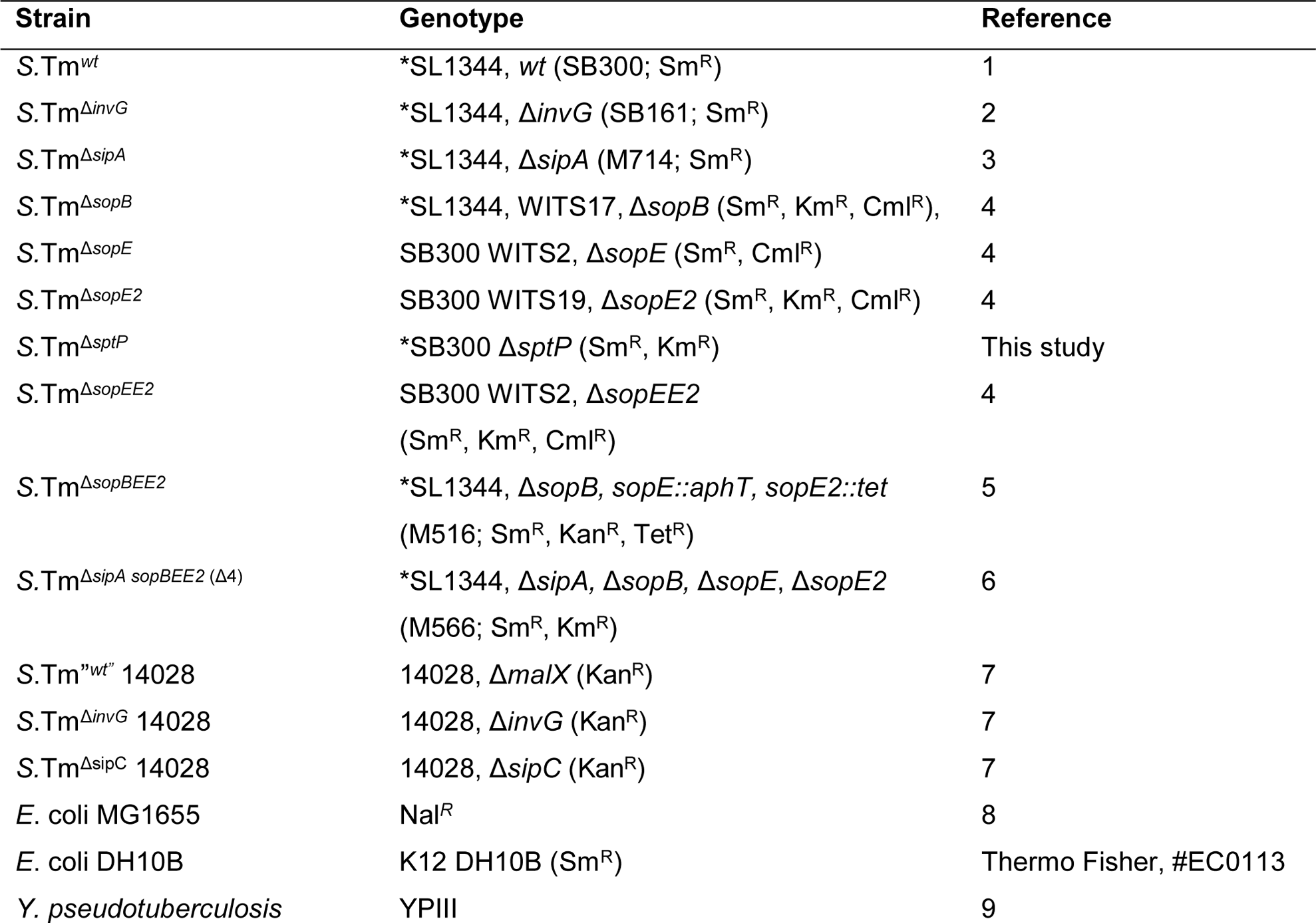
Bacterial strains and mutants used in this study. Indicated resistances as “Sm” for streptomycin, “Cml” for chloramphenicol, “Km” for kanamycin, “Tet” for tetracycline, and “Nal” for nalidixic acid. Strains marked with “*” prior the genotype were also used with the p*ssaG*-GFP reporter plasmid (S2 Table). The pFPV-mCherry plasmid (S2 Table) was used together with SL1344 *S*.Tm^wt^.

**S2 Table.** Plasmids used in this study. Plasmid Reference.

| Plasmid | Reference |
| --- | --- |
| pFPV-mCherry | 10 |
| *pssaG*-GFPmut2 high copy | 11 |

**S3 Table.**
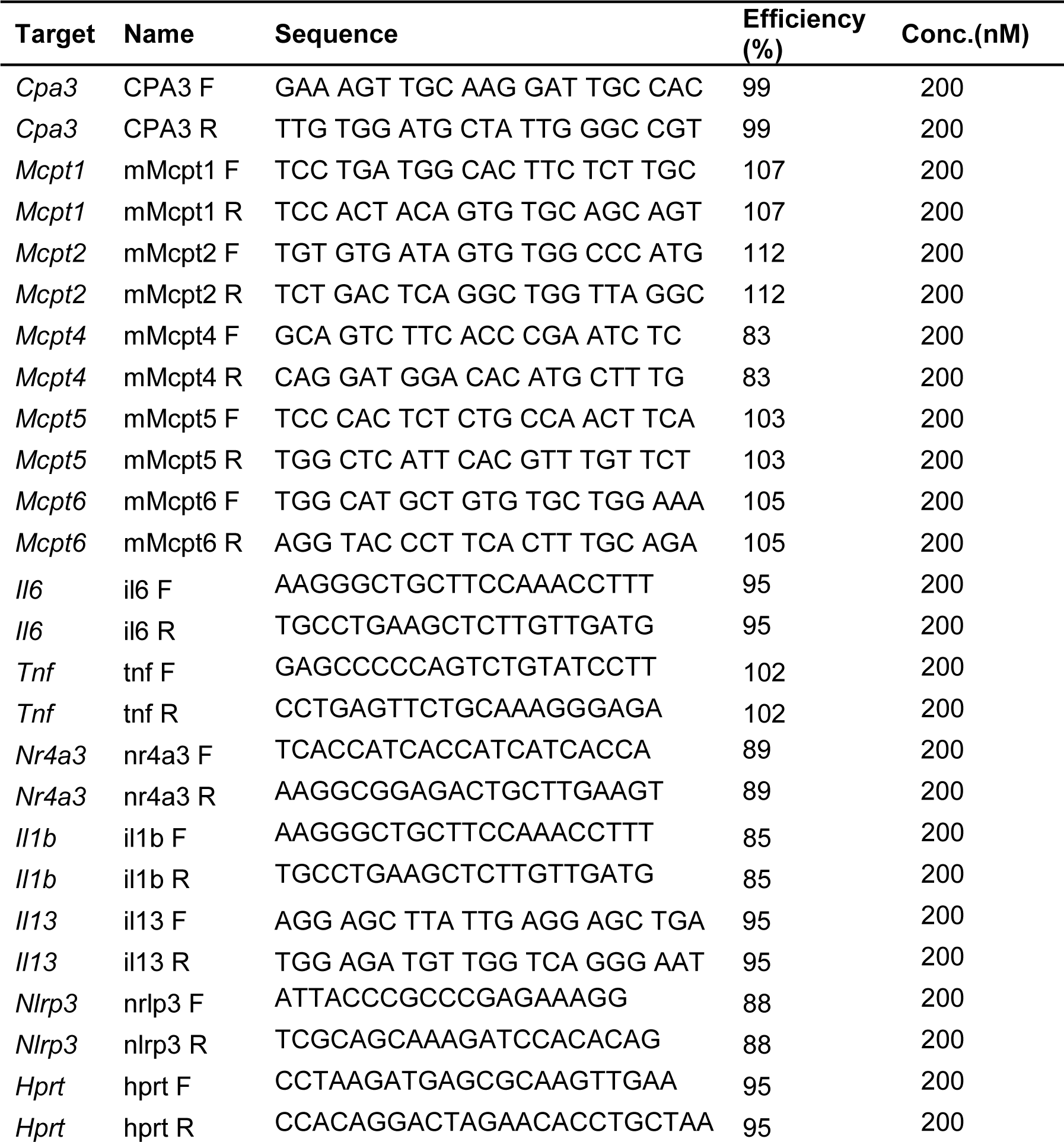
Primers used for RT-qPCR in this study.

**S4 Table.** Primary antibodies used in this study, with unique identifiers (RRID).

| Antigen | RRID | Conjugate | Host | Catalogue no | Clone | Provider | Dilution |
| --- | --- | --- | --- | --- | --- | --- | --- |
| *Salmonella* O Antiserum Factor 5 | NA | NA | Rabbit | BD-226601 | Polyclonal | Difco/MicLev | 1:250 |
| Akt | AB_329827 | NA | Rabbit | 9272 | Polyclonal | Cell Signaling | 1:1000 |
| Phospho-Akt (Ser473) | AB_2315049 | NA | Rabbit | 4060 | D9E | Cell Signaling | 1:2000 |
| CD16/32 | AB_394657 | NA | Rat | 553142 | 2.4G2 | BD Biosciences | 1:1000 |
| TLR4 | AB_469944 | Alexa Fluor 488 | Mouse | 53-9041-82 | UT41 | Invitrogen | 1:100 |
| Mouse IgG1 Isotype Control | NA | Alexa Fluor 488 | Mouse | MG120 | Polyclonal | Invitrogen | 1:100 |
| CD45 | AB_1645208 | Alexa Fluor 700 | Rat | 560510 | 30-F11 | BD Biosciences | 1:100 |
| CD11b | AB_468714 | PE-Cy5 | Rat | 15-0112-82 | M1/70 | eBioscience | 1:100 |
| CD11c | AB_469590 | PE-Cy7 | Armenian hamster | 25-0114-82 | N418 | eBioscience | 1:100 |
| F4/80 | AB_465923 | PE | Rat | 12-4801-82 | BMB | Invitrogen | 1:100 |
| CD45 | AB_470499 | NA | Rat | ab25386 | I3/2.3 | AbCam | 1:50 |
| CD18 | AB_396701 | NA | Rat | 557437 | M18/2 | BD Biosciences | 1:50 |

